# Dissection of the microRNA transcriptomes of CD4^+^ T cell subsets in autoimmune inflammation identifies novel regulators of disease pathogenesis

**DOI:** 10.1101/2025.06.23.661017

**Authors:** Carolina Cunha, Paula Vargas Romero, Daniel Inácio, Ana Teresa Pais, Catarina Pelicano, Marina Costa, Sofia Mensurado, Natacha Gonçalves-Sousa, Pedro H. Papotto, Daniel Neves, Daniel Sobral, Francisco Enguita, Bruno Silva-Santos, Anita Q. Gomes

**Author notes:** Corresponding authors (A.Q.G.) or (B.S.-S.). These authors contributed equally to this work (Bruno Silva-Santos, Anita Q. Gomes).

## Abstract

MicroRNAs (miRNAs) are key regulators of CD4^+^ T cell differentiation, but how they contribute to the course of an autoimmune disease *in vivo* remains poorly studied. Given the known roles in autoimmunity of pro-inflammatory T helper 1 (Th)1 and Th17 cells, and anti-inflammatory Foxp3^+^ regulatory cells, we established a triple reporter mouse for *Ifng*, *Il17* and *Foxp3,* and subjected it to experimental autoimmune encephalomyelitis (EAE) to characterize the miRNomes of the corresponding CD4^+^ T cell subsets. We identified 110 miRNAs differentially expressed between the pro-inflammatory (Th1 and Th17 cells) and the Treg cell subsets. Among these, we found novel functions for miR-122-5p and miR-1247 as regulators of Th17 cell proliferation and Th1 cell differentiation, thus impacting the course or severity of EAE, respectively. Importantly, their expression patterns suggest miR-122-5p and miR-1247 act as peripheral brakes to CD4^+^ T cell pathogenicity that are subverted in the inflamed central nervous system.

## Introduction

CD4^+^ T cells can differentiate into multiple functional subsets through networks of cytokines and transcription factors (TFs) that dictate lineage specific transcriptional programs. Interferon-γ (IFN-γ)-producing Thelper 1 (Th1) and interleukin (IL)-17A-producing Th17 cells are two of the key effector CD4^+^ T cell populations that orchestrate complementary immune responses to major pathogens, such as viruses and intracellular bacteria (Th1), or fungi and extracellular bacteria (Th17). Critically, these effector CD4^+^ T cells are also involved in the development of chronic inflammatory and autoimmune diseases such as type I diabetes, rheumatoid arthritis, psoriasis, colitis or multiple sclerosis (MS) (1, 2). On the other hand, CD4^+^ T cells can also differentiate, either in the thymus or in the periphery, into anti-inflammatory regulatory T cells (Treg) that are able to curtail the function of effector cells (Teff) (2, 3), and whose relevance is highlighted by the very severe autoimmune manifestations of Treg-defective IPEX patients and *Foxp3* mutant mice (4).

More specifically in MS, effector Th1 and Th17 CD4^+^ T cells contribute to disease pathogenesis after differentiation in the periphery and infiltration into the central nervous system (CNS), causing local inflammation and demyelination (5, 6). With regard to Th1 cells, genetic polymorphisms of IFN-γ have been associated with susceptibility to MS, and increased T cell-derived IFN-γ levels preceded disease exacerbation and correlated with active lesions *in vivo*, and killed oligodentrocytes *in vitro* (reviewed in 7). Th17 cells, on the other hand, activate microglia and recruit neutrophils, macrophages, and lymphocytes (8) through synergy of IL-17A with other proinflammatory cytokines such as IL-6, IL-23, IL-1β and GM-CSF (9,10). Critically, myelin oligodendrocyte glycoprotein (MOG)-specific Th1 and Th17 cells were shown to induce experimental autoimmune encephalomyelitis (EAE), the well-established mouse model of MS, upon adoptive transfer (6, 11, 12).

Immunosuppressive Tregs also play a role in MS by reducing the function of effector cells, often prevailing in remitting phases of the disease (13). In fact, when Treg cells were administered therapeutically in the EAE model, there was a reduction in disease severity, which coincided with limited immune cell infiltration, principally Th17 cells, into the CNS (14). These results reinforce the concept that the balance between Teff and Treg subsets is a major determinant of the outcome of the local inflammatory/ autoimmune response (2, 3). As such, it is important to identify molecular mechanisms that control Teff or Treg cells across their transition from the periphery to the target CNS.

MicroRNAs (miRNAs) have emerged as key players in the fine-tuning of gene expression at the post-transcriptional level. They are highly conserved small (∼22 nucleotide-long) non-coding RNAs that bind to the 3’UTR, and less frequently to the coding sequence (CDS), of target mRNAs to induce either degradation or translation repression (15, 16). miRNAs are known to regulate CD4^+^ T cell differentiation; most strikingly, the ablation of (all) miRNAs in T cells caused impaired T cell proliferation and survival, reduced Treg numbers, and increased production of IFN-γ and IL-17A by CD4^+^ T cells, leading to spontaneous lethal inflammatory disease (17, 18, 19, 20). However, only a few specific miRNAs have been identified as underlying such overt phenotypes. For example, miR-29 (21) and the miR-106-363 cluster (22) were shown to inhibit Th1 and Th17 differentiation, respectively, whereas miR-125a promoted Treg activity (23). In the EAE/ MS setting, miR-326 associated with Th17 cells and disease severity, and *in vivo* miR-326 silencing reduced Th17 numbers and EAE pathogenesis (24). Moreover, MS-enriched miR-92 limited Treg induction while supporting Th17 responses, and its inhibition ameliorated EAE (25).

Building on these foundations, we set out to characterize the global miRNomes of Th1, Th17 and Treg cells in EAE, with the ultimate goal of identifying novel miRNA regulators of CD4^+^ T cell functions *in vivo*. In order to gain resolution compared to previous studies, where candidates were selected based on the analysis of total CD4^+^ T cells of EAE mice or MS patients (compared to controls), we established a triple reporter mouse for IFN-γ (YFP^+^), IL-17A (GFP^+^) and Foxp3 (hCD2^+^) to isolate the corresponding CD4^+^ T cell subsets upon EAE induction. This allowed us to dissect their distinct miRNomes, and to identify new miRNAs, miR-122-5p and miR-1247-5p, that respectively regulate Th17 and Th1 cells, and thus impact EAE disease onset or severity *in vivo*.

## Results

### Identification of the miRNomes of *in vivo*-generated Th1, Th17 and Treg cells

In order to identify miRNA repertoires of CD4^+^ T cell subsets under inflammatory conditions *in vivo*, we established a triple reporter mouse for IFN-γ (YFP^+^), IL-17A (GFP^+^) and Foxp3 (hCD2^+^) (triple *Foxp3*^hCD2^.*Ifng*^YFP^.*Il17a*^GFP^ reporter mice **Fig. 1a**), by crossing previously described single reporter lines (26, 27, 28). As the response is triggered in peripheral lymphoid organs, we isolated these populations by FACS from the spleen and lymph nodes of EAE-induced triple *Foxp3*^hCD2^.*Ifng*^YFP^.*Il17a*^GFP^ reporter mice at the peak-plateau phase of the disease, and extracted RNA to perform both small RNA- and messenger-RNA sequencing (small RNA-seq and mRNA-seq, respectively) (**Fig. 1b, Fig. S1a**). Both datasets showed that the isolated YFP^+^, GFP^+^ and hCD2^+^ CD4^+^ T cell populations segregated unambiguously on a principal component analysis (**Fig. S1b and c, respectively**), and importantly, differentially expressed the signature genes of each corresponding CD4^+^ T cell subset (**Fig. S1d**), thus attesting the reliability and reproducibility of our experimental system. The small RNA-seq data revealed clearly distinct miRNomes, with 110 miRNAs differentially expressed between Teff and Treg cell populations *in vivo* (**Fig. 1c** and **d, Table S1**). These data are a valuable resource for the community, from which we decided to zoom in on miRNAs specifically enriched or depleted in one of the CD4^+^ T cell subsets. Interestingly, within this candidate list of differentially expressed miRNAs, only nine were specifically up- or down-regulated in one subset (i.e. Th1, Th17 or Treg) compared with the other two subsets (**Fig. 1e**): miR-1247-5p was up-regulated in Th1 cells; miR-126a-5p, miR-5108 and miR-122-5p were up-regulated in Th17 cells, miR-211-5p, miR-15b-5p, miR-151-3p and miR-467a-5p were up-regulated in Treg cells; and miR-125a-5p was down-regulated in Th17 cells. We performed validation studies by RT-qPCR analysis of independent Th1, Th17 and Treg samples isolated by FACS from the spleen/lymph nodes of triple reporter mice with EAE. These confirmed the expression profiles of six of the candidate miRNAs (**Fig. 1f**), with the exceptions of miR-151-3p, miR-5108 and miR-467a-5p. Notably, two of the validated miRNAs, miR-15b-5p and miR-125a-5p, have already been thoroughly studied and, consistent with our expression data, shown to be involved in Treg or Th17 differentiation and/or function (29, 30, 23, 30, 31). Thus, for subsequent “proof-of-concept” functional assays, and since the effector cell populations are the major drivers of EAE, we focused on the candidates displaying higher fold change expression levels in either Th1 or Th17 cells, miR-1247-5p and miR-122-5p, respectively (**Fig. 1e and f**).

**Fig. 1.**
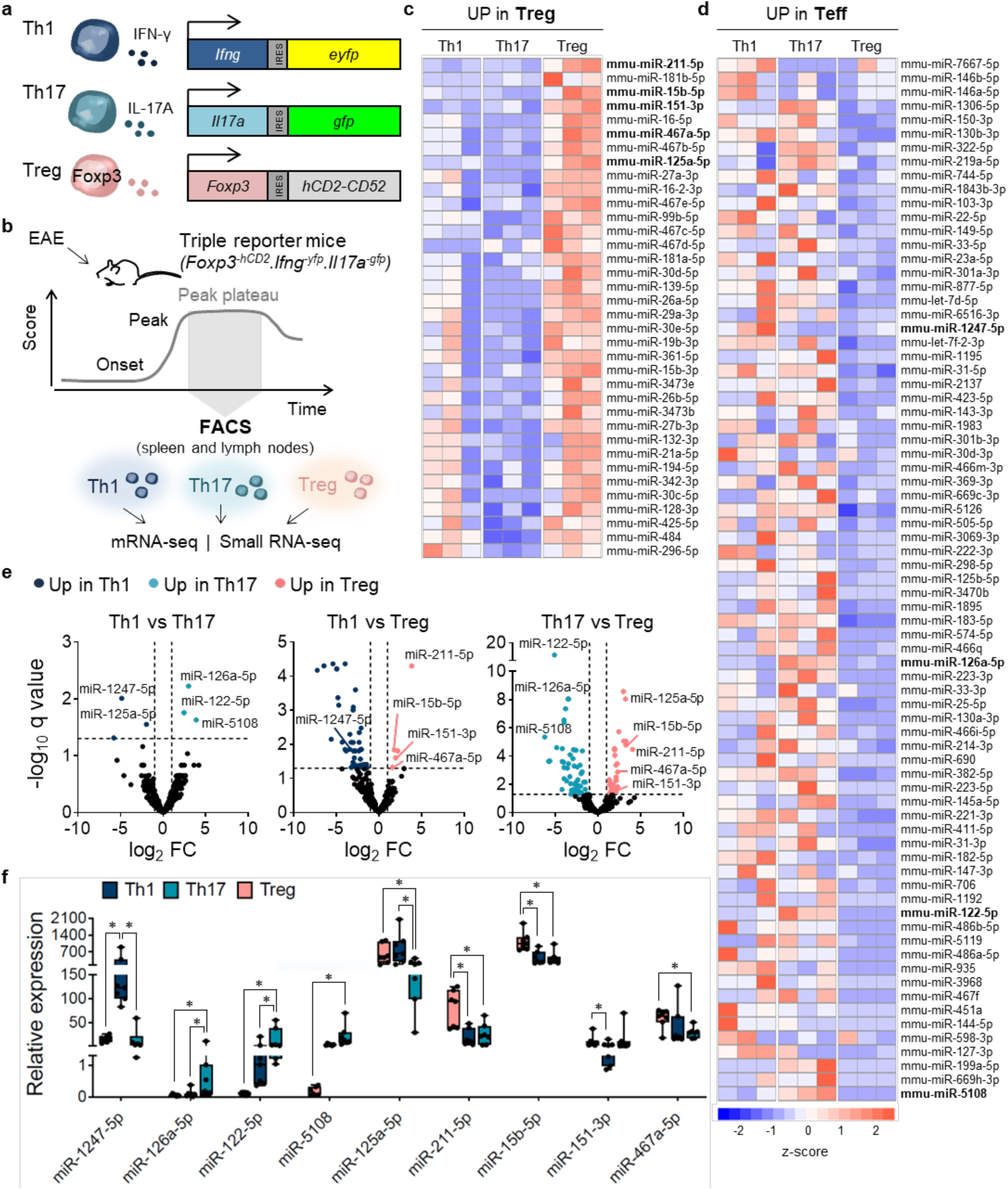
Identification of the miRNomes of Th1, Th17 and Treg cells from EAE-challenged *Foxp3*^hCD2^.*Ifng*^YFP^.*IL17a*^GFP^ reporter mice. (a) Schematic representation of the genomic alterations introduced in the triple *Foxp3-hCD2.Ifng-YFP.Il17a-GFP* reporter mouse strain with the respective CD4^+^ T cell subset. In the loci of each reporter gene, a sequence for an independently translated reporter protein was inserted downstream of the gene endogenous translational stop codon. (b) Experimental setup: triple *Foxp3-hCD2.Ifng-YFP.Il17a-GFP* reporter mice were subjected to EAE and at the peak plateau stage Treg, Th1 and Th17 cells were isolated from the spleen and lymph nodes by FACS, according with each reporter gene. After RNA isolation samples were subjected to both mRNA-seq and small RNA-seq. (c, d) Heatmaps depicting differentially expressed miRNAs between Treg and effector Th (up in Treg vs Th1 or Treg vs Th17 (c); versus up in Th1 vs Treg or Th17 vs Treg (d)). Colors indicate the direction and magnitude of calculated z-scores, with red representing higher expression of a miRNA in a given cell subset and blue, the opposite. (e) Volcano plots depicting differentially expressed miRNAs between Th1, Th17 and Treg subsets. miRNAs names are indicated when differentially expressed (log2FC > 1; FDR < 0.05) in two comparisons. (f) RT-qPCR analysis of the 9 candidate miRNAs’ expression in Treg, Th1 and Th17 cells isolated and pooled from the lymph nodes and spleen of triple Foxp3-hCD2.Ifng-YFP.Il17a-GFP reporter mice at EAE peak plateau. Box-and-whiskers indicate median, 25 and 75% percentiles, minimum and maximum expression levels of candidate miRNAs. Results are presented relative to miR-423-3p expression from 7 independent experiments. *p<0.05 (Wilcoxon test).

### Overexpression of miR-122-5p or miR-1247 directly impacts Th17 and Th1 cells *in vitro*

We inquired whether these miRNAs might have a cell-intrinsic role in Th17 (miR-122-5p) or Th1 (miR-1247) cells by performing overexpression experiments in *in vitro*-differentiated cells (from isolated naïve CD4^+^ T cells – **Fig. S2**). Upon transduction with retroviral GFP reporter vectors (RVs) encoding the native stem loop of miR-122 (RV-122) (**Fig. 2a**), we observed an upregulation of miR-122-5p levels in *in vitro*-differentiated Th17 cells (**Fig. S3a**). While no differences were observed in the frequency of IL-17A^+^ (**Fig. 2 b and c**), Th17 cells overexpressing miR-122 had a significantly lower proliferation rate when compared with control cells (RV-empty), as observed by the frequency of proliferating GFP^+^ T cells upon CTV labeling as well as their division index (**Fig. 2d-f**). Notably, miR-122 overexpression did not impact Th1 or Treg cell differentiation *in vitro* (**Fig. S3b and c**). These results suggest that miR-122-5p negatively regulates Th17 cell function by limiting their proliferative capacity.

**Fig. 2.**
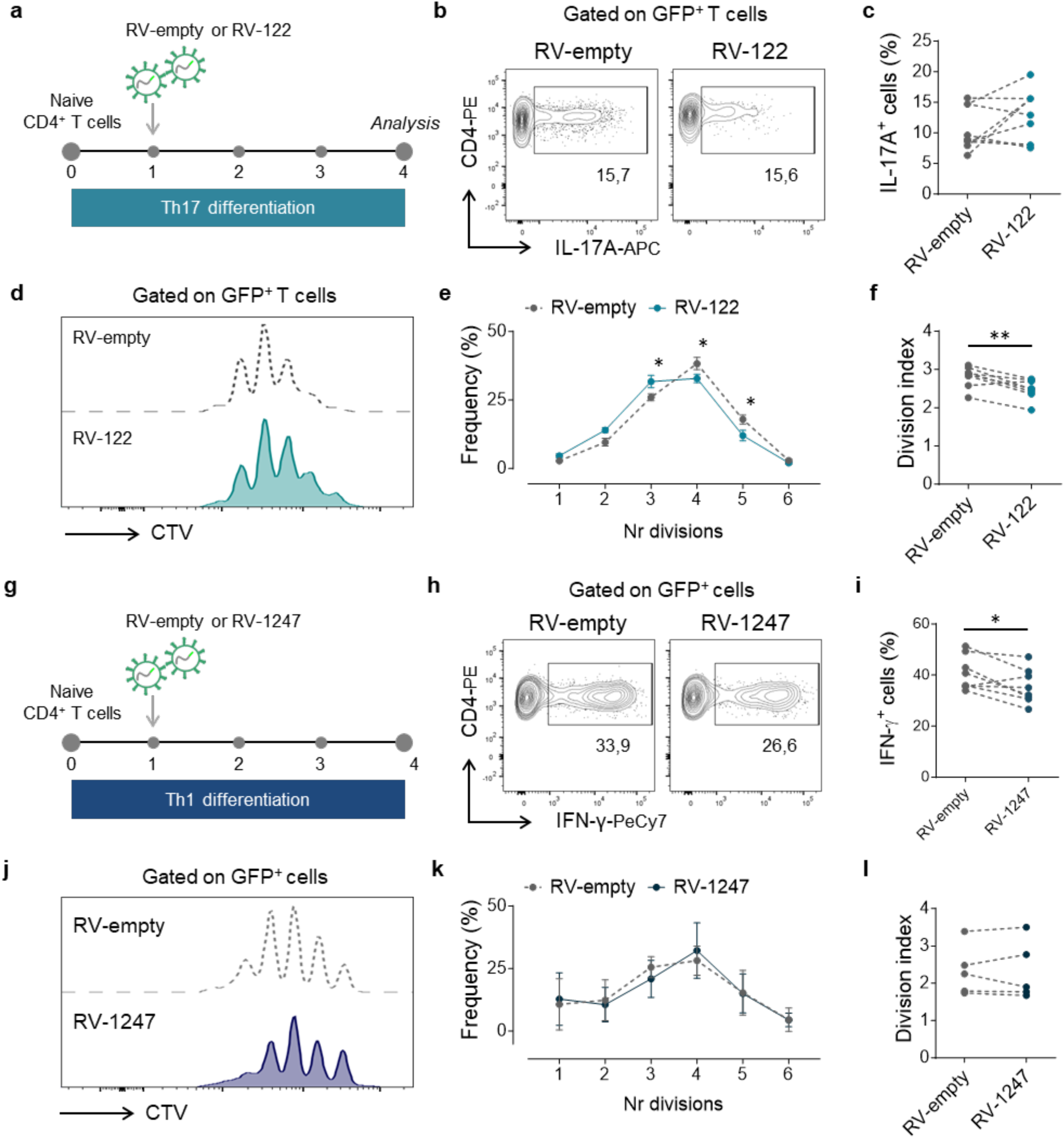
miR-122-5p and miR-1247 intrinsically impact Th17 cell proliferation and Th1 cell function. (a) Experimental setup: naive CD4^+^ T cells were sorted from WT mice and differentiated *in vitro* towards Th17 cells for 4 days. At day 1, cells were transduced with retroviral vectors encoding the precursor forms of miR-122 and differentiation was analysed by flow cytometry. Empty vector was used as control. (b) Representative flow cytometry plots of IL-17A^+^ cells in retrovirally transduced Th17 cells expressing a control vector (RV-empty) or miR-122 (RV-122) and frequency of IL-17A^+^ cells in retrovirally transduced Th17 cells expressing a control vector (RV-empty) or miR-122 (RV-122) from 5 independent experiments (c). Dotted lines link cells from the same mouse. (d) Representative histogram of proliferating CTV^+^ cells in retrovirally transduced Th17 cells expressing a control vector (RV-empty) or miR-122 (RV-122). (e) Frequency of proliferating CTV^+^ cells in retrovirally transduced Th17 cells expressing a control vector (RV-empty) or miR-122 (RV-122). Results are mean ± SD from 2 independent experiments (n = 4). *p<0.05 and **p<0.01 (two-way ANOVA followed by Sidak’s multiple comparisons test). (f) Division index of retrovirally transduced Th17 cells expressing a control vector (RV-empty) or miR-122 (RV-122). Dotted lines link cells from the same mouse. *p<0.05 (Paired t-test). (g) Experimental setup: naive CD4^+^ T cells were sorted from WT mice and differentiated in vitro towards Th1 cells for 4 days. At day 1, cells were transduced with retroviral vectors encoding the precursor forms of miR-1247 and differentiation was analysed by flow cytometry. Empty vector was used as control. (h) Representative flow cytometry plots of IFN-γ^+^ cells in retrovirally transduced Th1 cells expressing a control vector (RV-empty) or miR-1247 (RV-1247) and frequency of IFN-γ^+^ cells in retrovirally transduced Th1 cells expressing a control vector (RV-empty) or miR-122 (RV-122) from 5 independent experiments (i). Dotted lines link cells from the same mouse. *p<0.05 (Paired t-test). (j) Representative histogram of proliferating CTV^+^ cells in retrovirally transduced Th1 cells expressing a control vector (RV-empty) or miR-1247 (RV-1247). (k) Frequency of proliferating CTV^+^ cells in retrovirally transduced Th1 cells expressing a control vector (RV-empty) or miR-1247 (RV-1247). Results are mean ± SD from 3 independent experiments (n = 5). (l) Division index of retrovirally transduced Th1 cells expressing a control vector (RV-empty) or miR-1247 (RV-1247). Dotted lines link cells from the same mouse.

In what refers to miR-1247-5p, its overexpression with retroviral vectors in differentiating Th1 cells (**Fig. 2g, Fig. S3d**) led to a decreased frequency of IFN-γ^+^ CD4^+^ T cells (**Fig. 2h and i**), but did not affect cell proliferation (**Fig. 2 j-l**), nor Treg or Th17 *in vitro*-polarization (**Fig. S3c and e**). These results indicate that miR-1247-5p selectively down-regulates IFN-γ production in Th1 cells.

### miR-122-5p is a peripheral brake to Th17 pathogenicity that is overruled in the CNS

We next aimed at dissecting the molecular cues that regulate miR-122-5p and miR-1247-5p expression. For this, we sorted naïve CD4^+^ T cells from the spleen and lymph nodes of WT mice and cultured them in the presence of different cytokines involved in Th17 or Th1 cell differentiation, respectively. Starting with the Th17-biased miR-122-5p, its expression reached maximal levels in the presence of IL-6 and TGF-β, but decreased when stimulated with IL-1β and especially with IL-23 (**Fig. 3a**). IL-6 and TGF-β are the key cytokines involved in the polarization of naïve CD4^+^ T cells into the Th17 subset, while IL-1β and IL-23 are critical for lineage maintenance and the acquisition of a pathogenic phenotype (27). Thus, our data suggests that miR-122 might limit the pathogenicity of Th17 cells but is downregulated in a strong (IL-23/ IL-1β-rich) inflammatory environment. To build on this premise *in vivo*, we characterized miR-122-5p expression levels in CD4^+^ T cell subsets isolated from the CNS of triple *Foxp3*^hCD2^.*Ifng*^YFP^.*Il17a*^GFP^ reporter mice challenged with EAE. Interestingly, our data showed that miR-122-5p was markedly downregulated in cells isolated from the CNS compared with peripheral Th17 cells (**Fig. 3b**). To further provide biological context to these observations, we performed mRNA-seq in Th17 cells isolated from both sites, i.e. spleen/lymph nodes *vs* CNS. This showed that genes associated with a pathogenic phenotype – such as those encoding key cytokines as IFN-γ and GM-CSF (*Ifng* and *Csf2*), chemokines (*Ccl4* and *Cxcl3*) and TFs (*Tbx21* and *Runx1*) - were all increased in CNS-derived Th17 cells when compared to peripheral Th17 cells (**Fig. 3c**) (27). These results link miR-122 expression dynamics with the pathophysiology of EAE, with its downregulation in the CNS associating with enhanced pathogenicity of Th17 cells.

**Fig. 3.**
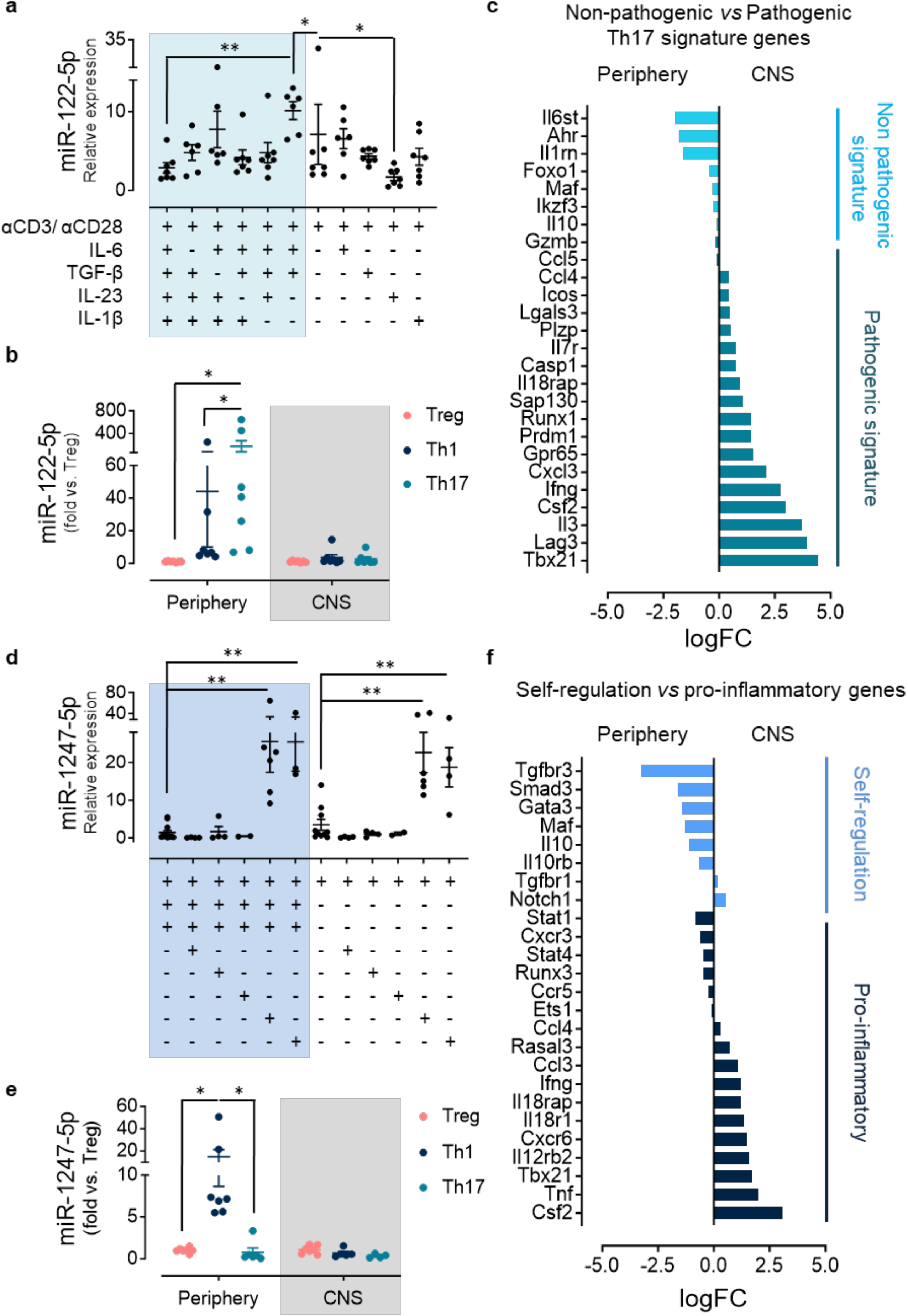
miR-122 and miR-1247 act as brakes on pro-inflammatory Th17/Th1 phenotypes that are subverted in the CNS. **(a)** RT-qPCR analysis of miR-122 expression in CD4^+^ T cells cultured *in vitro* for 4 days in the presence of indicated cytokines and antibodies. Results are presented relative to miR-423-3p expression and represent mean ± SD (n=6-7). *p<0.05 and **p<0.01 (non parametric Mann-Whitney test). **(b)** RT-qPCR analysis of miR-122 expression in Treg, Th1 and Th17 cells isolated and pooled from the periphery (lymph nodes and spleen) and the CNS of triple Foxp3-hCD2.Ifng-YFP.Il17a-GFP reporter mice at EAE peak plateau stage. Results are presented as fold vs. Treg in each specific tissue (ddct) and represent mean ± SD (n=7). miR-423-3p was used as reference gene. *p<0.05 (Wilcoxon test). **(c)** Gene expression determined by RNA-seq of pathogenic vs non pathogenic signature genes in Th17 cells isolated and pooled from the periphery (lymph nodes and spleen) and the CNS of triple Foxp3-hCD2.Ifng-YFP.Il17a-GFP reporter mice at EAE peak plateau stage. Results are represented as mean of log fold change (logFC) between periphery and CNS Th17 cells. **(d)** RT-qPCR analysis of miR-1247 expression in CD4^+^ T cells cultured *in vitro* for 4 days in the presence of indicated cytokines and antibodies. Results are presented relative to miR-423-3p expression and represent mean ± SD (n=2-12). *p<0.05 and p<0.01 (non parametric Mann-Whitney test). **(e)** RT-qPCR analysis of miR-1247 expression in Treg, Th1 and Th17 cells isolated and pooled from the periphery (lymph nodes and spleen) and the CNS of triple Foxp3-hCD2.Ifng-YFP.Il17a-GFP reporter mice at EAE peak plateau stage. Results are presented as fold vs. Treg in each specific tissue (ddct) and represent mean ± SD (n=7). miR-423-3p was used as reference gene. *p<0.05 (Wilcoxon test). **(f)** Gene expression determined by RNA-seq of self-regulation vs proinflammatory genes in Th1 cells isolated and pooled from the periphery (lymph nodes and spleen) and the CNS of triple Foxp3-hCD2.Ifng-YFP.Il17a-GFP reporter mice at EAE peak plateau stage. Results are represented as mean of log fold change (logFC) between periphery and CNS Th1 cells.

### miR-1247 limits Th1 pathogenicity in the periphery but is subverted in the CNS

We performed similar studies for miR-1247, the Th1-biased miRNA that downregulated IFN-γ production *in vitro* (**Fig. 2g**). Although miR-1247-5p was upregulated on *in vivo*-generated Th1 cells (**Fig. 1e and f**), its expression levels were low in Th1 cells differentiated *in vitro* in the presence of IL-12 and anti-IL-4 antibody (**Fig. 3d**). We also tested IL-18, IL-15 and IFN-γ, since IL-18 and IL-15 can promote IFN-γ production in the presence of IL-12 (32); and IFN-γ regulates its own production in an autocrine loop (33). However, none of these cytokines was able to induce miR-1247-5p expression in Th1 cells (**Fig. 3d**). We then reasoned that, as miR-1247-5p limits IFN-γ production by Th1 cells, its expression might be controlled by anti-inflammatory cytokines. Indeed, we found that TGF-β and IL-10 were the key inducers of miR-1247-5p expression *in vitro*, even in the presence of IL-12 (**Fig. 3d**). Moreover, when we challenged our triple reporter mice with EAE, we observed that, similarly to miR-122-5p (**Fig. 3b**), the expression levels of miR-1247-5p collapsed in Th1 cells isolated from the CNS when compared to peripheral Th1 cells (**Fig. 3e**). Furthermore, upon mRNA-seq analysis of Th1 cells isolated from both sites (spleen/lymph nodes *vs* CNS), we established a CNS-specific pro-inflammatory signature, including the signature Th1 cytokine (*Ifng*) and TF (*Tbx21*), as well as chemokines (*Ccl3* and *Ccl4*) and receptors (*Il12rb2*, *Il18r1*, *Il18rap*), which contrasted with a set of “self-regulatory genes”, including *Il10*, that were upregulated in Th1 cells from the secondary lymphoid organs (**Fig. 3f**). These data suggest that miR-1247 is part of an anti-inflammatory (TGF-β/ IL-10-mediated) regulatory loop that is subverted once Th1 cells invade the CNS and shut down its expression.

### Identification of putative mRNA targets of miR-122-5p in Th17 cells

Given that miRNAs exert their functions by suppressing mRNAs, and each miRNA can impact the expression of many mRNAs (34), we next aimed at identifying the repertoires of mRNAs directly regulated by miR-122-5p and miR-1247-5p. We devised a strategy based on differential Ago2-immunoprecipitation followed by RNA-seq (Ago2 RIP-seq) in lineage-specific cells overexpressing each candidate miRNA (upon retroviral transduction) versus control cells (mock-transduced) (**Fig. S4a**). Briefly, naïve CD4^+^ T cells were sorted and activated under polarizing conditions (Th1 or Th17 as appropriate), transduced at day 1 with pre-miR RV particles (as used above for the overexpression experiments) and further incubated for 36h, when GFP^+^ T cells were sorted and subjected to UV-cross linking and Ago2 immunoprecipitation with high affinity antibody (**Fig. S4a**).

Starting with miR-122-5p, RT-qPCR analysis after the Ago-IP confirmed its enrichment in the Argonaute complex of retrovirally miR-122 transduced Th17 cells (RV-122) relative to the empty vector control (RV-empty), whereas other miRNAs, such as miR-126a-5p or miR15b-5p, showed similar levels between RV-122 and RV-empty treated cells (**Fig. S4b**). The results of the differential Ago2 RIP-seq in Th17 cells identified 587 mRNAs overexpressed in miR-122-5p-transduced Th17 cells, of which 339 (57,8%) had predicted miR-122-5p binding sites (**Fig. S4c**). From these, as proof-of-concept, we inspected our mRNA-seq data for those upregulated in the CNS compared to peripheral Th17 cells, based on the previous observation that miR-122-5p had the opposite expression pattern (**Fig. 3b**). This retrieved 27 mRNAs (FC>1,5, p<0,05) (**Fig. 4a**), including 16 whose function was of relevance for Th17 cell biology and/or proliferation (**Table S2**). Based on literature curation, we further analyzed the expression levels of 5 genes (*Pttglip*, *Nudt4*, *Il18rap*, *Spred2* and *Sap130*) in retrovirally transduced Th17 cells with the native stem loop of miR-122 (RV-122), and observed that *Nudt4*, *Il18rap* and *Sap130* were specifically downregulated in miR-122 overexpressing (versus control) cells, further reinforcing their potential to be functional targets of miR-122 in the context of Th17 cells in EAE (**Fig. 4b**). Finally, to biochemically validate these 3 mRNAs as direct miR-122-5p targets, we performed luciferase assays. The three putative targets have predicted miR-122-5p binding sites in the 3’UTR of *Nudt4* and *Il18rap* and the CDS of *Sap130* (**Fig. 4c**). To determine whether miR-122-5p was directly targeting these mRNAs, we designed reporter constructs in a pmirGLO Dual-luciferase miRNA target expression vector for the 3′UTRs of *Nudt4* and *Il18rap* and the CDS of *Sap130*. We additionally included a negative control vector containing a 3′UTR without miR-122-5p binding sites and a positive control containing the 3′UTR of *Cpeb1*, a previously validated miR-122-5p target (35). We transiently transfected these constructs into human embryonic kidney (HEK) 293 T cells together with an expression plasmid for either miR-122-5p or an unrelated miRNA. Co-transfection of miR-122-5p (but not an unrelated miRNA) showed a significant repression of luciferase activity compared with control for the three mRNA targets (**Fig. 4c**). Mutations in the miR-122-5p binding sites of *Nudt4*, *Il18rap* and *Sap130* led to significant recoveries of luciferase levels, attesting specificity (**Fig. 4c**). Together, these data suggest that miR-122-5p targets *Nudt4*, *Il18rap* and *Sap130* in Th17 cells. Furthermore, it provides a much larger Ago2 RIP-seq data set that constitutes a resource for those interested in assessing miR-122-5p in future studies on T cell differentiation.

**Fig. 4.**
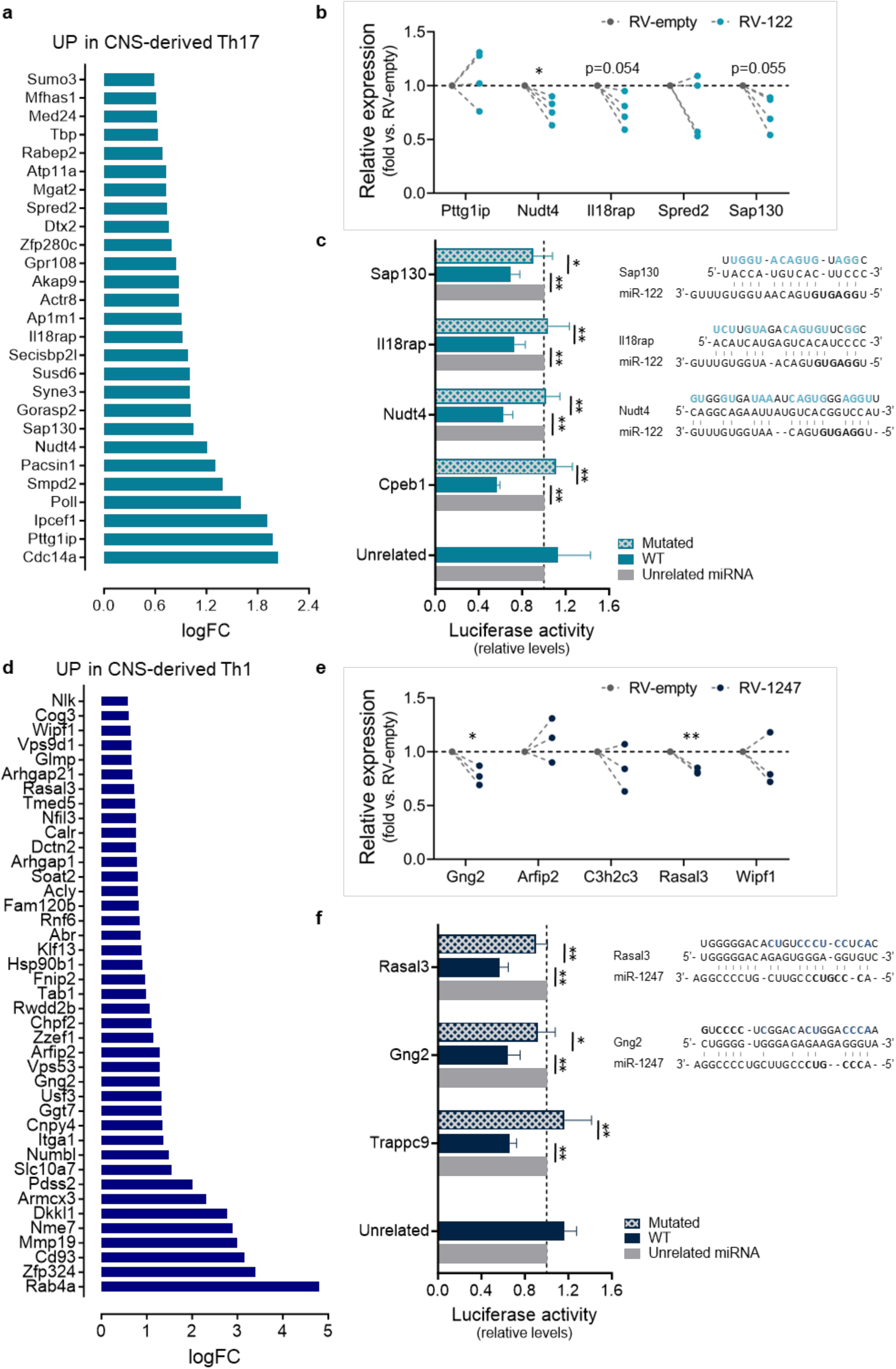
Identification of putative mRNA targets of miR-122-5p and miR-1247-5p in Th17 and Th1 cells. (a) Gene expression determined by RNA-seq of candidate miR-122-5p mRNA targets that are upregulated in Th17 cells isolated from the CNS of triple Foxp3-hCD2.Ifng-YFP.Il17a-GFP reporter mice at EAE peak plateau stage when compared to Th17 isolated from the periphery (lymph nodes and spleen) of the same mice (FC>1,5; p<0.05). Results are represented as mean of log fold change (logFC) between periphery- and CNS-derived Th17 cells. (b) RT-qPCR analysis of candidate miR-122-5p mRNA targets in retrovirally transduced Th17 cells expressing a control vector (RV-empty) or miR-122 (RV-122). Results are presented as fold vs. RV-empty (n=4). Grey dashed lines link cells from the same mouse. Black dashed line represents the baseline expression level of controls. *p<0.05 (paired students t-test). (c) Luciferase activity in HEK293T cells co-transfected with a pmirGLO Dual-Luciferase miRNA target expression vector containing either the WT or mutated miR-122-5p-binding sites of Sap130, Il18rap and Nudt4 mRNAs plus a vector containing miR-122-5p or an unrelated miRNA. An unrelated construct (without miR-122-5p binding sites) and a positive construct (Cpeb1) were included. Schematic representation of miR-122-5p putative binding sites of in the miR response elements of Sap130, Il18rap and Nudt4 mRNAs are included. Results are presented as fold vs. unrelated miRNA and represent mean + SD (n=5). *p<0.05 and **p<0.01 (paired students t-test). (d) Gene expression determined by RNA-seq of candidate miR-1247-5p mRNA targets that are upregulated in Th1 cells isolated from the CNS of triple Foxp3-hCD2.Ifng-YFP.Il17a-GFP reporter mice at EAE peak plateau stage when compared to Th1 isolated from the periphery (lymph nodes and spleen) of the same mice (FC>1,5; p<0.05). Results are represented as mean of log fold change (logFC) between periphery- and CNS-derived Th1 cells. (e) RT-qPCR analysis of candidate miR-1247-5p mRNA targets in retrovirally transduced Th1 cells expressing a control vector (RV-empty) or miR-1247 (RV-1247). Results are presented as fold vs. RV-empty (n=3). Grey dashed lines link cells from the same mouse. Black dashed line represents the baseline expression level of controls. *p<0.05 (paired students t-test). (f) Luciferase activity in HEK293T cells co-transfected with a pmirGLO Dual-Luciferase miRNA target expression vector containing either the WT or mutated miR-1247-5p-binding sites of Rasal3 and Gng2 mRNAs plus a vector containing miR-1247-5p or an unrelated miRNA. An unrelated construct (without miR-1247-5p binding sites) and a positive construct (Trappc9) were included. Schematic representation of miR-1247-5p putative binding sites of in the miR response elements of Rasal3 and Gng2 mRNAs are included. Results are presented as fold vs. unrelated miRNA and represent mean + SD (n=5). *p<0.05 and **p<0.01 (paired students t-test).

### Identification of putative mRNA targets of miR-1247-5p in Th1 cells

We employed the same experimental approach to identify miR-1247-5p targets in the context of Th1 cell differentiation. The differential Ago2 RIP-seq (**Fig. S4a**) showed that the RNA-induced silencing complex (RISC) of miR-1247-overexpressing Th1 cells (as confirmed by RT-qPCR, **Fig. S4b**) was enriched (compared to mock-transduced Th1 cells) in 644 mRNAs, of which 423 (65,7%) had predicted binding sites for miR-1247-5p (**Fig. S4c**). Upon examination of our mRNA-seq data for Th1 cells, we found 41 mRNAs upregulated (FC>1,5 and p>0,05) in the CNS (**Fig. 4d**), of which 18 had relevant functions for Th1 cell biology (**Table S2**). Based on literature curation, we selected 5 genes - *Gng2*, *Arfip2*, *C2h2c3*, *Rasal3* and *Wipf1-* to further test *in vitro*, i.e. upon retroviral transduction of the native stem loop of miR-1247 in Th1 cells (Fig. 4e). Of these, *Gng2* and *Rasal3* were validated as miR-1247-5p targets both *in vitro*, as showed by their downregulation upon miR-1247-5p overexpression (**Fig. 4e**), and in luciferase assays demonstrating direct binding of *Gng2* and *Rasal3* using the same strategy as described above for miR-122-5p target genes (**Fig. 4f**). More specifically, the 3’UTR regions of both wild-type and mutant *Gng2* and *Rasal3* (**Fig. 4f**) were introduced into a pmirGLO Dual-luciferase miRNA target expression vector and co-expressed with either miR-1247-5p or a control unrelated miRNA. These assays showed a decrease in luminescence in the presence of the miRNA that was abolished when the 3’UTR target regions were mutated (**Fig. 4f**). Thus, our results indicate that miR-1247-5p targets *Gng2* and *Rasal3*. Moreover, they provide the community with a comprehensive Ago2 RIP-seq data set for additional studies on miR-1247-5p in the context of effector T cell differentiation.

### miR-122-5p inhibition increases the frequency of IL-17A^+^ cells and anticipates EAE onset

Next, to assess if the candidate miRNAs play a role in EAE pathophysiology, we treated mice challenged with EAE with synthetic antagomiRs and evaluated their impact on disease course and on the frequency of Teff and Treg subsets in the lymph nodes of antagomiR-treated mice at the end of the experiment. As depicted in **Fig. 5a**, we injected the antagomiRs of the candidate miRNAs at days 7, 9 and 11 after immunization, which corresponds to a preclinical stage (onset of symptoms usually occurs around day 10). Focusing first on miR-122-5p antagomiR treatment - which efficiently reduced miR-122-5p expression levels (**Fig. S5a**) - we observed an anticipation of EAE disease onset as mice started to develop symptoms and lose weight earlier than those treated with the control antagomiR (**Fig. 5b**). Importantly, all mice that were subjected to miR-122-5p antagomiR injections were symptomatic already at day 12 post immunization (p.i.), whereas the control group needed an additional 5 days (day 17 p.i.) to present symptoms, thus entering the peak plateau stage later on (**Fig. S5b**).

**Fig. 5.**
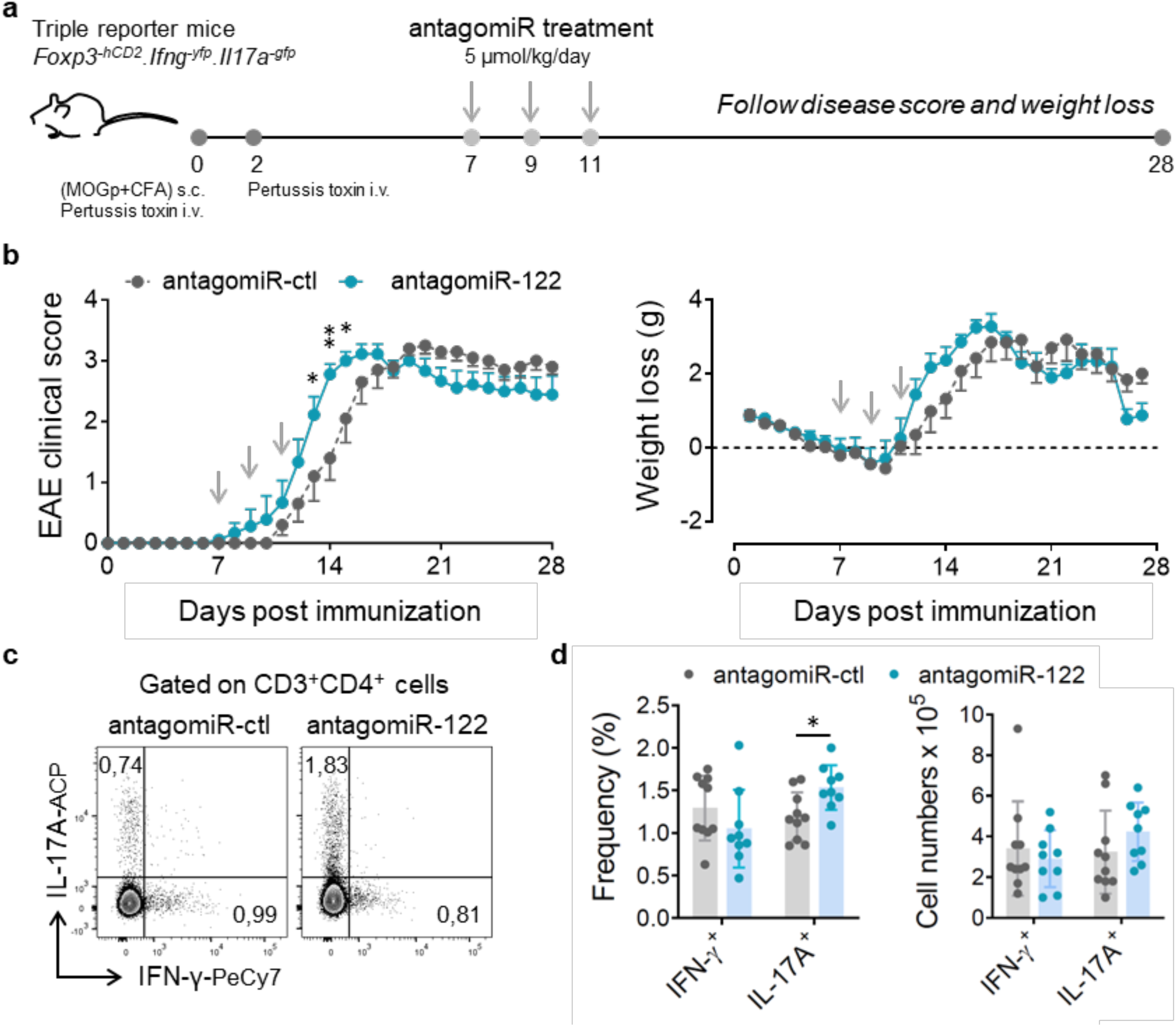
miR-122-5p inhibition antecipates EAE onset. **(a)** Experimental setup: triple *Foxp3-hCD2.Ifng-YFP.Il17a-GFP* reporter mice were immunized s.c. in both flanks with 100 µg of myelin oligodendrocyte glicoprotein (MOG)^35-55^ peptide emulsified in CFA solution (day 0). On the day of immunization and 2 days later, mice received a retro orbital injection of 200 ng of pertussis toxin. At days 7, 9 and 11, mice received 5 µmol/kg of specific antagomiRs (retro orbital injection). AntagomiR for cel-miR-67-3p was used as control. **(b)** Mice treated with control and miR-122-5p antagomiRs were scored for EAE clinical signs and weighed daily for 28 days. Results are mean ± SD from 3 independent experiments (n = 9-11). *p<0.05 and **p<0.01 (two-way ANOVA followed by Sidak’s multiple comparisons test). **(c)** Representative flow cytometry plots of IFN-γ^+^ and IL-17A^+^ CD4^+^ T cells in the cervical and lumbar lymph nodes of triple *Foxp3-hCD2.Ifng-YFP.Il17a-GFP* reporter mice with EAE and treated with control and miR-122-5p antagomiRs. **(d)** Frequency and absolute numbers of IL-17A^+^ CD4^+^ T cells in the cervical and lumbar lymph nodes of triple *Foxp3-hCD2.Ifng-YFP.Il17a-GFP* reporter mice with EAE and treated with control and miR-122-5p antagomiRs. Results are mean + SD from 3 independent experiments (n = 9-11). Each dot represents an individual mouse. *p<0.05 (non-parametric Mann-Whitney test).

Analysis of cervical and lumbar LNs (cLNs) showed no differences in the total CD4^+^ T cell frequency **Fig. S5c**), but an increased percentage of IL-17A^+^ CD4^+^ T cells in mice subjected to miR-122-5p antagomir treatment when compared with controls (**Fig. 5c and d**), consistent with the early onset of the disease. We found no differences in the frequency and total cell numbers of Th1 (IFN-γ^+^) and Treg (Foxp3^+^) cell populations in this case (**Fig. 5d**, **Fig. S5d and e**). Moreover, no differences were observed in the frequency of other immune cells between mice treated with miR-122-5p or control antagomiRs (**Fig. S5f**) based on the gating strategy highlighted in **Fig. S6a** (T cell populations) and **Fig. S6b** (other immune cell populations).

To assess the selectivity of these phenotypes, we studied the impact of an anti-miR-126a antagomiR. While this resulted in the downregulation of miR-126a (**Fig. S7a**), it had no impact neither on clinical scores nor on weight loss, since the specific antagomir-treated mice followed a similar disease course as the control group (**Fig. S7b**), and no differences were found on the frequency of IFN-γ^+^ and IL-17A^+^ nor Treg (Foxp3+) cell populations (**Fig. S7c**). These data highlight our finding that miR-122-5p inhibition selectively increases the frequency of Th17 cells and anticipates EAE onset.

### Upregulation of miR-1247 limits IFN-γ production by Th1 cells and reduces EAE severity

We next analysed the *in vivo* effects of miR-1247 manipulation. Unlike the previous antagomiRs (**Fig. S5a**), the miR-1247-5p construct led to increased levels of miR-1247-5p in various tissues (**Fig. S8a**), thus constituting an overexpression strategy. Consistent with this, we found decreased levels of previously validated miR-1247 targets, namely *Trappc9* and *Synj2bp*, in matching organs (**Fig. S8b).** While the progression of symptoms followed a similar course (**Fig. S9a**), we observed decreased EAE severity, as well as reduced weight loss, in mice overexpressing miR-1247-5p compared with the control group (**Fig. 6a**). Furthermore, this phenotype associated with a selective decrease in the numbers of IFN-γ^+^ CD4^+^ T cells (**Fig. 6b and c**), without affecting the total CD4^+^ T cell frequency (**Fig. S9b**), in the draining lymph nodes of miR-1247-5p-treated group. Of note, the frequency of other immune cells also did not change upon miR-1247-5p overexpression (**Fig. S9c**). Thus, the upregulation of miR-1247 expression restricts IFN-γ production by Th1 cells and reduces EAE severity.

**Fig. 6.**
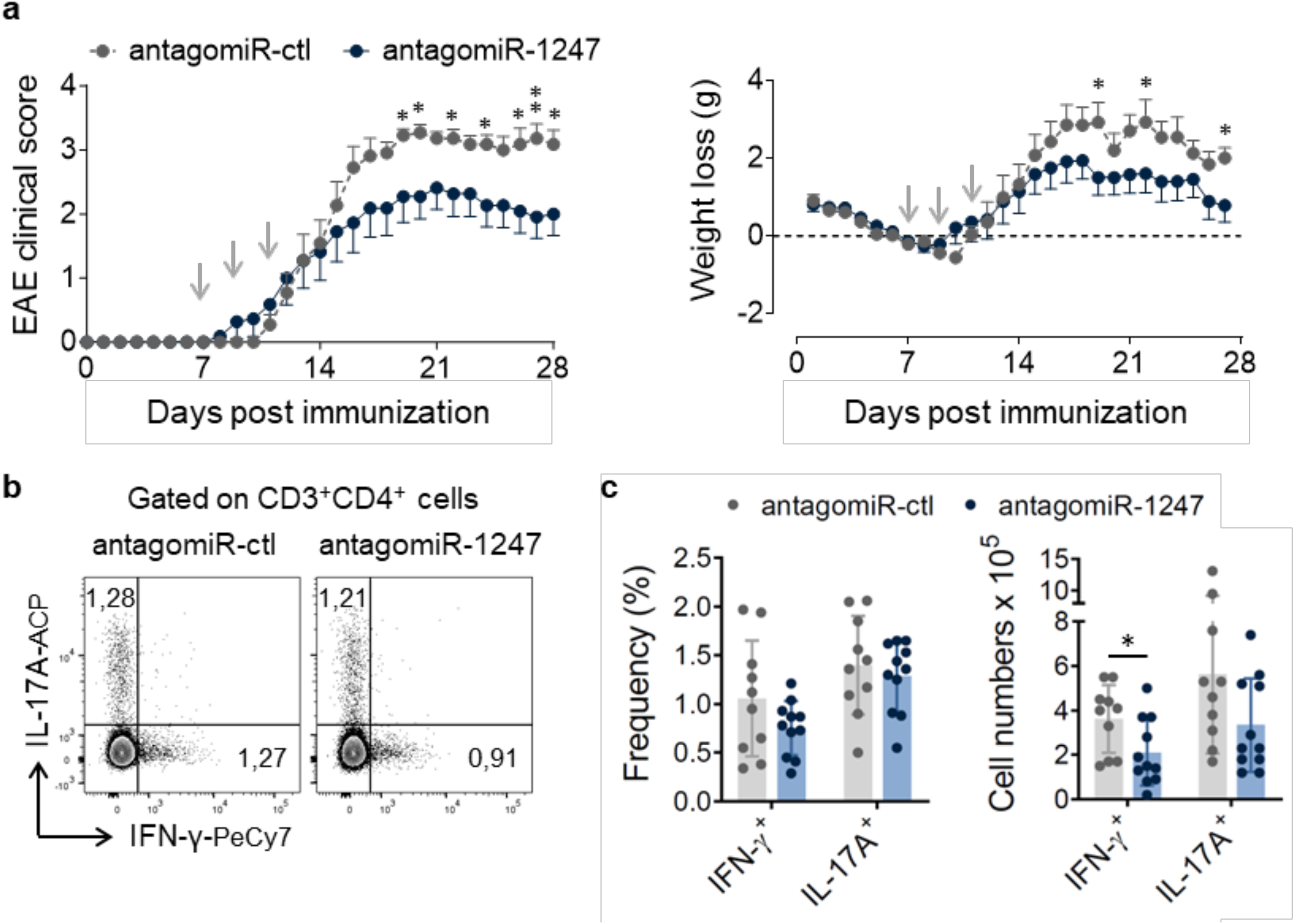
miR-1247-5p upregulation limits EAE severity. **(a)** Mice treated with control and miR-1247-5p antagomiRs were scored for EAE clinical signs and weighed daily for 28 days. Results are mean ± SD from 3 independent experiments (n = 10-11). *p<0.05 and **p<0.01 (two-way ANOVA followed by Sidak’s multiple comparisons test). **(b)** Representative flow cytometry plots of IFN-γ^+^ and IL-17A^+^ CD4^+^ T cells in the draining lymph nodes (inguinal, axillary and brachial) of triple Foxp3-hCD2.Ifng-YFP.Il17a-GFP reporter mice with EAE and treated with control and miR-1247-5p antagomiRs. **(c)** Frequency and absolute numbers of IFN-γ^+^ CD4^+^ T cells in the draining lymph nodes (inguinal, axillary and brachial) of triple Foxp3-hCD2.Ifng-YFP.Il17a-GFP reporter mice with EAE and treated with control and miR-1247-5p antagomiRs. Results are mean + SD from 3 independent experiments (n = 10-11). Each dot represents an individual mouse. *p<0.05 (non parametric Mann-Whitney test).

Collectively, our *in vivo* experiments have identified two novel miRNAs, miR-1247-5p and miR-122, that control Th1 and Th17 cell pathogenicity and thus impact the disease course of EAE.

## Discussion

This study aimed to provide novel insights on how miRNAs regulate CD4^+^ T cell subsets during an autoimmune response *in vivo*. Previous studies on the topic had focused on a limited set of miRNAs, such as miR-29^21^ and the miR-106-363 cluster (22) which were shown to inhibit Th1 and Th17 differentiation, respectively, by directly targeting either T-bet (and Eomes) or RORγt, TFs that directly regulate the production of IFN-γ or IL-17. Furthermore, several miRNAs had been shown to be either up-regulated – e.g. let-7e, miR-155, miR-223 and miR-326 – or down-regulated – e.g. miR-20b, miR-26a and miR-30a - in CD4^+^ Th cell subsets of MS patients or EAE-challenged mice, with some being functionally implicated in MS or EAE – for example, miR-326 upregulation stimulated the production of IL-17 in Th17 cells and enhanced pathogenesis (24) while miR-92 restricted Treg induction and supported Th17 responses, consistent with its elevated levels in CD4^+^ T cells of MS patients (25). Notably, two of our validated miRNAs, miR-15b-5p and miR-125a-5p, have already been shown to control Treg/Th17 cell function in EAE. Thus, miR-15b-5p, which was downregulated in EAE and MS, suppressed Th17 cells while increasing peripheral Treg differentiation (29, 30); and miR-125a-5p stabilized the immunoregulatory capacity of Treg cells, thereby impacting EAE severity (23, 31).

We now provide a global resource on miRNAs, as well as mRNAs, expressed on Th1, Th17 and Treg cell subsets from the peripheral lymphoid organs and the CNS (for mRNA-seq) of EAE-challenged mice, which can be exploited in further studies. The functional analyses of miR-122-5p and miR-1247 as “proof-of-concept” for our resource provided a key conceptual insight: they constitute molecular brakes to pro-inflammatory Th1 and Th17 cells that are subverted in the CNS. Thus, miR-122-5p and miR-1247 were strikingly downregulated in respectively Th17 and Th1 cells from the CNS compared with their lymphoid organ counterparts, and this was accompanied by an upregulation of pro-inflammatory/ pathogenic signatures (like *Ifng*, *Tbx21* or *Csf2*, the later encoding GM-CSF) at the mRNA level. We further identified the molecular cues that regulate the dynamic expression of these two miRNAs. For Th17-biased miR-122, we found it to be induced by IL-6 and TGF-β, but downregulated by IL-1β and especially IL-23. This is particularly interesting because while IL-6 and TGF-β are the key cytokines promoting Th17 cell polarization from naïve CD4^+^ T cells, IL-1β and IL-23 are critical for Th17 cell maintenance, expansion and, most importantly, the acquisition of a pathogenic phenotype in the target organ (27). As for the Th1-biased miR-1247-5p, we identified TGF-β and IL-10 as the key inducers of its expression, even in the presence of IL-12. Given that TGF-β induces the expression of IL-10 in Th1 cells to reduce their encephalogenicity (36, 37), we conceptualize miR-1247 as part of this self-regulatory mechanism to control excessive IFN-γ levels in Th1 cells.

To gain molecular insight on miR-122-5p and miR-1247 functions, we aimed to characterize direct mRNA targets relevant to Th17 or Th1 cell biology, respectively. In fact, mRNA target identification arguably remains the greatest challenge in miRNA biology, as it is now clear that each miRNA can fine-tune the expression of hundreds of mRNAs in any given cell type (34). To tackle this problem, and in line with the resource nature of our study, we used differential Ago2 RIP-seq on *in vitro*-polarized CD4^+^ T cells: by overexpressing the candidate miRNA, we promoted its binding to target mRNAs, thus enriching them in Ago2 complexes that are UV-cross linked and immunoprecipitated, followed by RNA-seq. This resulted in large Ago2 RIP-seq datasets that can be used by the community for other studies on these miRNAs in effector CD4^+^ T cells. Here we zoomed in on candidate mRNAs targets using the following criteria: 1) having predicted binding sites for the candidate miRNA; 2) being upregulated in the CNS compared to secondary lymphoid organs, thus inversely correlating with the miRNA expression; 3) being downregulated upon retroviral overexpression of the miRNA *in vitro*; 4) directly binding to the mRNA sequence as indicated by luciferase assays.

Using this pipeline, we identified *Nudt4*, *Il18rap* and *Sap130* as putative validated targets of miR-122-5p in Th17 cells. Nudt4 is a hydrolase that catalyses the m^7^G tRNA modification that promotes the translation of cell cycle mRNAs (38), consistent with the Th17 cell proliferation phenotypes upon miR-122-5p modulation. On the other hand, polymorphisms in the IL18R1-IL18RAP locus, an IL-18 receptor-associated and responsive protein, have been associated with autoimmune disease susceptibility and IL18RAP was shown to be associated with pathogenic Th17 signatures (39, 40, 41). Moreover, IL-18 has been reported to play a similar role to IL-1β in the promotion of IL-17 production and autoimmunity (41). Finally, Sap130 function remains more enigmatic, but it is known to associate with Sin3A (42), which is essential for IL-17 production (43). Collectively, the identification of these 3 direct targets provide new biological context to miR-122-5p-mediated regulation of Th17 cells.

With regard to miR-1247, our analysis pinpointed *Gng2* and *Rasal3* as direct targets in Th1 cells. Gng2, G-protein gamma subunit 2, acts as a tumor suppressor gene by inhibiting Akt and Erk activity (44). In turn, Erk1-deficient mice exhibit increased Th1 cell differentiation and earlier EAE onset than wild-type controls (45). As for Rasal3, Ras GTPase activating protein-like 3, it is a member of the GTPase-activating proteins family required for survival of naïve and activated T cells (46). Importantly, both Rasal3 and its partner GTPase, Rac2 (47) have been shown to promote T-cell production of IFN-γ (48, 49). This previous knowledge on Gng2 and Rasal3 nicely fits our proposed model where miR-1247-mediated targeting impacts Th1 responses in EAE. Overall, by meeting the criteria stated above, these validated mRNA targets offer mechanistic insight into the functions of miR-122-5p and miR-1247 in regulating the pathogenicity of T helper cell subsets in EAE.

Notably, our functional analyses demonstrated the pathophysiological impact of miR-122 and miR-1247 on EAE onset and severity, which associated with intrinsic effects on Th17 cell proliferation and Th1 cell differentiation, respectively. As both miRNAs suppress these pro-inflammatory responses, our findings highlight the potential of modulating specific miRNAs to control the pathogenic immune populations implicated in autoimmune diseases like MS.

A limitation of our current study is that it does not address the role of miRNAs in CD4^+^ T cell plasticity during EAE (50). Th17 cells, in particular, are known to transdifferentiate into Th1 cells under the IL-23-rich inflammatory environment of EAE (51), but we opted for a snapshot of the effector CD4^+^ T cell subsets at the peak of EAE using dynamic reporter mice, rather than fate mapping (50). We also did not address the impact of our miRNA candidates on the production of GM-CSF, which has been shown by Becher and colleagues to initiate autoimmune neuroinflammation (52), particularly with its secretion by Th1 and Th17 cells being sufficient to induce EAE (5, 53). Conversely, the key contributions of this study are the global resource on miRNAs expressed in Th1, Th17 and Treg cells at the peak of EAE disease severity; the identification of two new miRNA determinants of Th1/ Th17 cell function, their repertoires of putative mRNA targets in those cells, the intricate regulation by a network of cytokines and their expression dynamics *in vivo* across lymphoid organs and the CNS, which open new avenues to explore in both mice and humans, with the ultimate goal of devising miRNA-based immunotherapies for MS.

## Resource availability

### Lead contact

Anita Quintal Gomes

### Materials availability

All unique/stable reagents generated in this study are available from the lead contact without restriction.

### Data and code availability

CD4^+^ T cell RNA-seq data are available from the Sequence Read Archive (SRA) under the accession code PRJNA1067547 (https://www.ncbi.nlm.nih.gov/bioproject/PRJNA1067547). All other data that support the findings of this study are available in the article and Supplementary Information.

### Author contributions

C.C., A.Q.G., and B.S.-S. designed the research and wrote the paper; C.C., P.V.R., D.I., A.T.P., C.P., D.S., M.C., S.M., N.G.-S., P.H.P., F.E. and A.Q.G. performed experiments and analyzed the data; B.S.-S. and A.Q.G. supervised the research.

## Material and Methods

### Mice

All mice used were females 6 to 12 weeks of age. C57Bl/6J (B6) WT mice were purchased from Charles River Laboratories. Triple *Foxp3-hCD2.Ifng-YFP.Il17a-GFP* reporter mice were generated and bred in house by crossing the following single reporter strains: B6.Foxp3hCD2 (http://www.informatics.jax.org/allele/MGI:4950682) (54), Great (Ifng-YFP) (http://www.informatics.jax.org/allele/MGI:5317415) (26), and IL17-GFP (http://www.informatics.jax.org/allele/MGI:5426354) (27). B6.Foxp3hCD2 and Great mice were obtained from the Jackson Laboratory and Il17a-GFP mice were from Biocytogen, LLC. Mice were bred and maintained in the specific pathogen–free animal facilities of Gulbenkian Institute for Molecular Medicine (Lisbon, Portugal). All experiments involving animals were approved by the Ethics Animal Welfare Body of Gulbenkian Institute for Molecular Medicine (ORBEA-GIMM), set up in accordance with Portuguese law (article 34 of Decreto-Lei 113/2013, transposed from the European Directive 2010/63/EU), and submitted to the local competent authority Direcção-Geral de Alimentação e Veterinária (DGAV) for authorization. Euthanasia was performed by CO2 inhalation and anesthesia was performed by isofluorane inhalation.

### EAE induction and scoring

Triple *Foxp3-hCD2.Ifng-YFP.Il17a-GFP* reporter mice were immunized s.c. in both flanks with 100 µg of myelin oligodendrocyte glycoprotein (MOG) 35–55 peptide (MEVGWYRSPFSRVVHLYRNGK) (Eurogentec S.A.) emulsified in CFA solution (4 mg/ml of heat-inactivated M. tuberculosis in IFA) (Difco Laboratories). On the day of immunization and 2 days after, mice received 200 ng pertussis toxin (PTx) (List Biological Laboratories) in 100 µl PBS i.v.. Mice were weighed daily and scored for EAE clinical signs. In brief, the score system ranged from 0 to 5, with 0.5 increments, being score 0 attributed to animals with no clinical signs of EAE and five representative of death. Score 1 consisted in limp tail; score 2 consisted in limp tail together with hind legs weakness; score 3 consisted in complete limb paralysis; and finally, score 4 consisted in complete hind leg paralysis and partial front paralysis. Mice were euthanized if they lose 20% of body weight or if they scored 4 for 2 consecutive days.

### AntagomiR treatment

AntagomiRs were custom synthesized according to (55) (Merck). Antagomir sequences are as follows: antagomiR-1247-5p -5’-U*C*CGGGGACGAACGGGACG*G*G*U*-chol; antagomiR-122-5p - 5’- C*A*AACACCAUUGUCACACU*C*C*A*-chol; antagomiR-126a-5p - 5’- C*G*CGUACCAAAAGUAAUA*A*U*G*-chol; AntagomiR for cel-miR-67-3p was used as negative control: antagomiR-ctl - 5’-U*C*UACUCUUUCUAGGAGGUUG*U*G*A*-chol. All ribonucleotides were 2′-O-methyl modified and (*) represents a phosphorothioate modification of the backbone. A cholesterol molecule was added at the 3′-end of the oligonucleotides. Lyophilized antagomiRs were resuspended in RNase-free water at the desired concentration at room temperature for 30 min with slight shaking. *Foxp3-hCD2.Ifng-YFP.Il17a-GFP* reporter mice with EAE were treated with 5 µg/kg specific antagomiRs at days 7, 9 and 11 after immunization, which corresponds to a preclinical stage (onset of symptoms occurs around day 10) through i.v. injection in the eye. Treatments were randomized per cage and EAE scoring was performed by a researcher blinded to each animal’s treatment group.

### Monoclonal antibodies

The following anti-mouse fluorescently labeled monoclonal antibodies (mAbs) were used (antigens and clones): CD3 (145.2C11), CD4 (GK1.5 and RM4-5), CD25 (PC61), CD44 (IM7), CD62L (DREG.55), IFN-γ (XMG1.2), IL-17A (eBio17B7), Foxp3 (FJK-16s), CD8 (53–6.7), TCRδ (GL3), F4/80 (BM8), NK1.1 (PK136), CD19 (6D5), Ly6G (1A8), Ly6C (HK1.4), CD11B (M1/70), CD11C (N418) and CD45 (30-F11). The anti-human CD2 (RPA-2.10) antibody was also used. Antibodies were purchased from BD Biosciences, eBiosciences, or BioLegend.

### Cell preparation, cell sorting, and flow cytometry and analysis

Cell suspensions were obtained from spleens, lymph nodes (cervical, axillary, brachial, inguinal and lumbar), and the central nervous system (brain and spinal cord). For the preparation of lymph nodes and spleen, tissues were filtered through a 70-µm cell strainer and erythrocytes from the spleen were osmotically lysed in red blood cell lysis buffer (BioLegend). For the preparation of brain and spinal cord, mice were perfused through the left cardiac ventricle with cold PBS. Brain and spinal cord were dissected and cut into pieces, and further digested with collagenase type IV (1.5 mg/ml; Roche) and DNase I (0.10 mg/ml) (Sigma-Aldrich) in RPMI 1640 containing 5% fetal bovine serum (FBS) for 30 min, at 37°C. Mononuclear cells were isolated by passing the tissue through a 100-µm cell strainer, followed by a 40-70% Percoll (Sigma-Aldrich) gradient and 30-min centrifugation at 2400 rpm. Mononuclear cells were recovered from the interphase, resuspended, and used for further analysis.

For cell surface staining, single-cell suspensions were incubated in the presence of anti-CD16/CD32 (eBioscience) with saturating concentrations of combinations of the antibodies listed above for 15 min. For viability assessment, cells were stained in the presence of LIVE/DEAD Fixable Near-IR (Thermo Fisher Scientific) or Zombie Aqua (BioLegend), according with manufacturer’s instructions.

For intracellular cytokine staining or reporter analysis, cells were stimulated with PMA (phorbol 12-myristate 13-acetate) (50 ng/ml) and ionomycin (1 µg/ml), in the presence of Brefeldin A (10 µg/ml) (all from Sigma) for 3-3,5h at 37°C. Cells were stained for the identified above cell surface markers, fixed for 30 min at 4°C with the Foxp3/Transcription Factor Staining Buffer set (eBioscience), and finally incubated for 45-60 min at 4°C with identified above intracellular antibodies in permeabilization buffer. Cells were analyzed using FACSFortessa (BD Biosciences) and FlowJo software (Tree Star).

For cell sorting, single cell suspensions were stained for cell surface markers as mentioned above and then electronically sorted on a FACSAria (BD Biosciences). Cell sorting based on reporter genes, included stimulation with PMA/ionomycin/Brefeldin A before the staining.

### RNA isolation, complementary DNA synthesis, and RT-qPCR

RNA was isolated from fluorescence-activated cell sorting (FACS)–sorted cell populations using an miRNeasy Mini kit (Qiagen). For total RNA, reverse transcription was performed with random oligonucleotides (Invitrogen) using Moloney murine leukemia virus reverse transcriptase (Promega). For miRNA, reverse transcription was performed with a Universal cDNA Synthesis kit II (Exiqon). For total RNA samples, relative quantification of specific complementary DNA (cDNA) species to endogenous references HPRT, β-2-microglobulin or β-actin was carried out using SYBR on a ViiA7 cycler (Applied Biosystems). Primers were either designed manually or by the Universal ProbeLibrary Assay Design Center (Roche), and their sequences are indicated in the **Table S3**. For miRNA samples, relative quantification of specific cDNA species to reference miR-423-3p was carried out using SYBR on a ViiA7 cycler (Applied Biosystems). The respective miRNA LNA primers were used (Exiqon). In both cases, relative quantification was performed using ddCT method as indicated in each section.

### RNA- and small-RNA-sequencing

Deep sequencing (small non-coding RNA and RNA-seq) was performed at the GeneCore facility of EMBL (http://www.genecore.embl.de/). For RNA-seq samples were processed with NEBNext Ultra II RNA Library kit for Illumina. Specifically, the protocol used was that for purified mRNA or rRNA depleted RNA, with an optimization of the following steps to accommodate the low input: Fragmentation time (4.1.3) 10 minutes, First strand synthesis step 2 (4.2.3) was increased to 50 minutes for a longer insert, Adaptor was diluted to 1:30 (4.5.4) and final PCR (4.8.3) was increased to 18 cycles. For small RNA-seq samples were processed with the NEBNext Small RNA Library Prep Set for Illumina.

### miRNA-Seq data analysis

Sequenced reads were processed using the CAP-miRSeq pipeline (56). Namely, reads were quality filtered and adaptors removed using cutadapt. Filtered reads were aligned to the mouse genome (GRCm38) using bowtie (57). miRNA prediction and quantification was performed using miRDeep2 (58), where known miRNAs were obtained from mirbase (version 21) (59). Differential expression analysis was performed using the edgeR R package (60). Namely, count data was normalized using the TMM normalization (61), and moderated bayesian statistics (62) was applied to obtain differentially expressed genes (FDR < 0.05 and [Log2FC] > 1) between conditions. Comparisons were performed between Treg and Th (i.e. combining Th17 and Th1), Treg and Th1, and Treg and Th17.

### RNA-Seq data analysis

Sequenced reads were quality filtered and adaptors removed using cutadapt. Filtered reads were aligned to the mouse genome (GRCm38) using Hisat2 (63). Gene count tables were obtained using featureCounts (64) against the mouse gene models (gencode M16).

### *In vitro* CD4^+^ T cell subset polarization

Naive CD4^+^ T cells (CD3^+^ CD4^+^ CD25^-^ CD62L^+^CD44^low^) were FACS-sorted from the LNs and spleen of B6 WT mice and differentiated under polarizing conditions for 4 days. For Th1 cell culture conditions, naive CD4^+^ T cells were incubated with plate-bound anti-CD3ε (145.2C11) and soluble anti-CD28 (37.51) antibodies (both at 2 μg/ml) in the presence of IL-12 (10 ng/ml) and neutralizing anti–IL-4 antibody (11B11) (5 μg/ml). For Treg cells, naive CD4^+^ T cells were incubated with plate-bound anti-CD3ε and soluble anti-CD28 (both at 2 μg/ml) in the presence of IL-2 (10 ng/ml) and TGF-β (2 ng/ml). For Th17 cell polarization, naive CD4^+^ T cells were incubated with plate-bound anti-CD3ε (1 μg/ml) and anti-CD28 (10 μg/ml) in the presence of IL-1β (10 ng/ml), IL-6 (20 ng/ml), IL-23 (20 ng/ml), TGF-β (2 ng/ml) and neutralizing anti–IFN-γ antibody (R4-6A2) (10 μg/ml). All cytokines were from PeproTech, except TGF-β and IL-23, which were from R&D systems.

### Retroviral overexpression of miR-122 and miR-1247

The retroviral constructs encoding either mmu-miR-122 and mmu-miR-1247 were generated by inserting the respective native pre-miRNA sequences flanked by about 200 bp into a modified pMig.IRES-GFP retroviral vector (Addgene #9044), as previously reported (65). For retroviral transduction, CD4^+^ T cells were sorted from LNs and spleen of C57BL/6J mice and polarized towards Th17, Th1 or Treg cells as mentioned above. At polarization day 1, cells were transduced with retrovirus encoding the precursor forms of miR-122 (RV-122), miR-1247 (RV-1247) or the control vector (RV-empty), in the presence with polybrene (8 µg/ml; Sigma-Aldrich). At day 4, differentiation was analysed by flow cytometry cells and gene expression by qPCR.

### Proliferation assay

For analysis of proliferation, naïve CD4^+^ T cells were stained with 1 mM of CellTrace Violet (Thermo Fisher Scientific) in PBS for 20 minutes at room temperature. Reactions were quenched by washing with ice-cold RPMI medium supplemented with 10% (v/v) FBS. CTV-labeled cells were then polarized towards Th1 or Th17 cells and, on the next day, retrovirally transduced with retrovirus encoding the precursor forms of miR-122 (RV-122), miR-1247 (RV-1247), respectively, or the control vector (RV-empty), as previously described. After 4 days, proliferation was assessed by flow cytometry.

### Differential argonaute 2 RNA immunoprecipitation followed by deep sequencing

Th17 and Th1 were retrovirally transduced with RV-control vector and RV-miR-122 or RV-miR-1247 vector, respectively. Seventy-two hours after transduction, GFP^+^ live cells were FACS-sorted and UV-cross-linked in Stratalinker 2400 (400 mJ cm−2). Cell lysates were pulled from independent experiments (∼1 million Th17 and ∼400 000 Th1 cells per replicate) and lysed on ice for 15 min before immunoprecipitation. Argonaute (Ago) protein immunoprecipitation was performed using Protein G Dynabeads (Invitrogen) and the 2A8 anti-Ago antibody (Millipore). Briefly, Dynabeads were coated with AffiniPure rabbit anti-mouse IgG antibody (Jackson ImmunoResearch) for 50 min at room temperature, and incubated for additional 4h at 4°C with the 2A8 anti-Ago antibody (both with spinning). After incubation, cell lysates were added to antibody-coated beads and incubated overnight at 4°C while spinning. Ago2-bound RNA was then partially digested using RNase T1 (ThermoFisher Scientific) and, after proteinase K treatment, RNA was isolated using phenol chloroform isoamyl (Fisher Scientific) extraction. Deep sequencing was performed as above mentioned.

### Ago2 RIP-seq data analysis

Raw Illumina sequence reads obtained by the pair-end sequencing strategy were quality-checked with the FastQC tool (66) (version 0.11.9; Babraham Bioinformatics, UK) and were filtered and trimmed, with Trimmomatic software (67). Filtered and trimmed sequence reads were aligned with the mouse genome (genome build GRCm38, release 102) using the STAR aligner (68) (version 2.7.10b). Differential gene expression among sample groups was analysed by the DESeq2 algorithm, implemented in the Nasqar platform for high-throughput sequencing data analysis and visualization. Predicted miRNA targets for mmu-miR-1247-5p and mmu-mir-122 were determined by the miRWalk2 platform.

### Dual luciferase reporter assay

Plasmid vectors pMig-miR-122-5p and pMig-miR-1247-5p that allow for the overexpression of miR-122-5p or mir-1247-5p, respectively, were generated as described above. The binding regions of miR-122-5p targets – Nudt4, Il18rap and Sap130 and of miR-1247-5p targets Gng2 and Rasal13 and respective regions with mutations in the predicted target sequences were directly synthesised into pmiRGLO vector (GeneCust Europe). Each luciferase reporter vector carrying one of the sequences described above was co-transfected with either the pMig-miR-122-5p or pMig-miR-1247-5p expression vectors or a control empty pMig vector into HEK293 T cells (ATCC CRL-3216) using X-tremeGENE HP DNA Transfection Reagent (Roche). After 48 h, firefly and Renilla luciferase activity were measured by using the Dual-Glo Luciferase Assay System (Promega). Renilla luciferase activity served as the internal control, and relative luciferase activity was normalized to empty pMirGlo and to empty pMig-IRES-GFP.

### Statistical analysis

The statistical significance of differences between two groups was assessed using non-parametric two-tailed Mann-Whitney test or, if both groups followed a normal distribution (tested by Shapiro-Wilk normality test), using two-tailed unpaired Student’s t-test. Wilcoxon-matched-pairs or paired t-test were used for paired samples, following normality testing as above. When more than two groups were compared, two-way ANOVA followed by Sidak’s post hoc test was performed. Unless otherwise indicated, individual values and mean are plotted, or mean ± SD. p < 0.05 was considered significant and is indicated on the figures.

## Acknowledgements

We thank Vladimir Benes and his team at European Molecular Biology Laboratory GeneCore facility for assistance with library preparation for both the RNA and small-RNA-sequencing; and the staff of the Flow Cytometry and Rodent facilities of GIMM for valuable technical assistance. This work was funded by the European Research Council (CoG_646701 to B.S.-S.); and by Fundação para a Ciência e Tecnologia (SFRH/BD/145352/2019 PhD grant to D. I.).

## Supporting information

**Fig S1.**
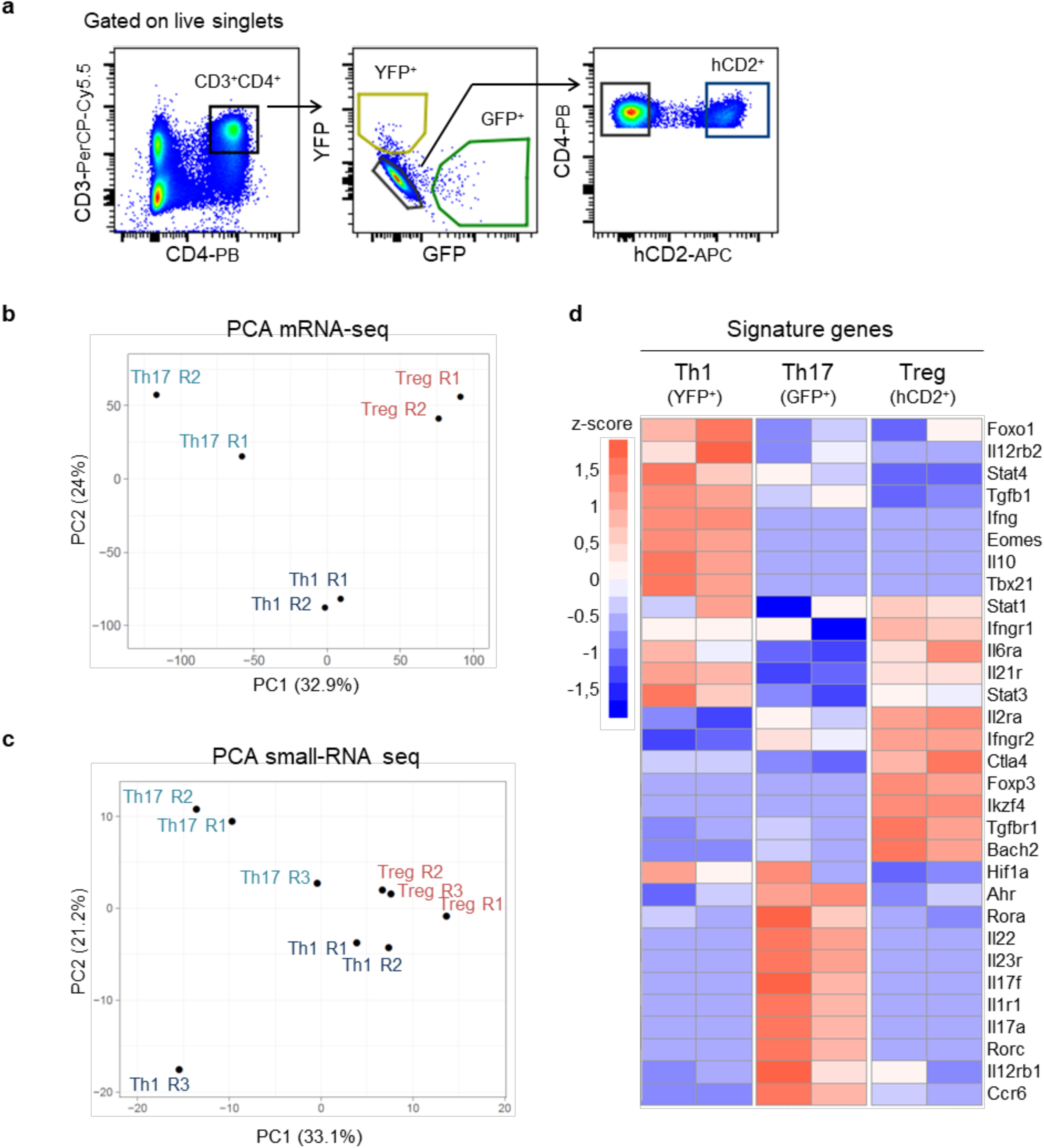
*In vivo* isolation of Th1, Th17 and Treg cell populations and their global characterization. (a) FACS gating strategy to isolate pure Th1, Th17 and Treg cells from EAE mice. Lymphocytes were selected based on a forward scatter (FSC)/side scatter (SSC) plot. After selection of live singlets, lymphocytes were gated on a CD4/CD3 plot to select CD4^+^ T cells. CD4^+^ lymphocytes were then gated on a GFP/YFP plot to sort Th1 cells (GFP^-^YFP^+^) and Th17 cells (GFP^+^YFP^-^). Finally, Treg cells were defined as GFP^-^YFP^-^hCD2^+^. (b-c) Principal component analysis (PCA) of the *in vivo*-generated Treg, Th1 and Th17 subsets subjected to RNA-seq. (c) and to small RNA-seq. (d) Heatmap depicting Treg, Th1 and Th17 signature genes that are enriched within, YFP^+^, GFP^+^ and hCD2^+^sorted populations, respectively. Colors indicate the direction and magnitude of calculated z-scores.

**Fig. S2.**
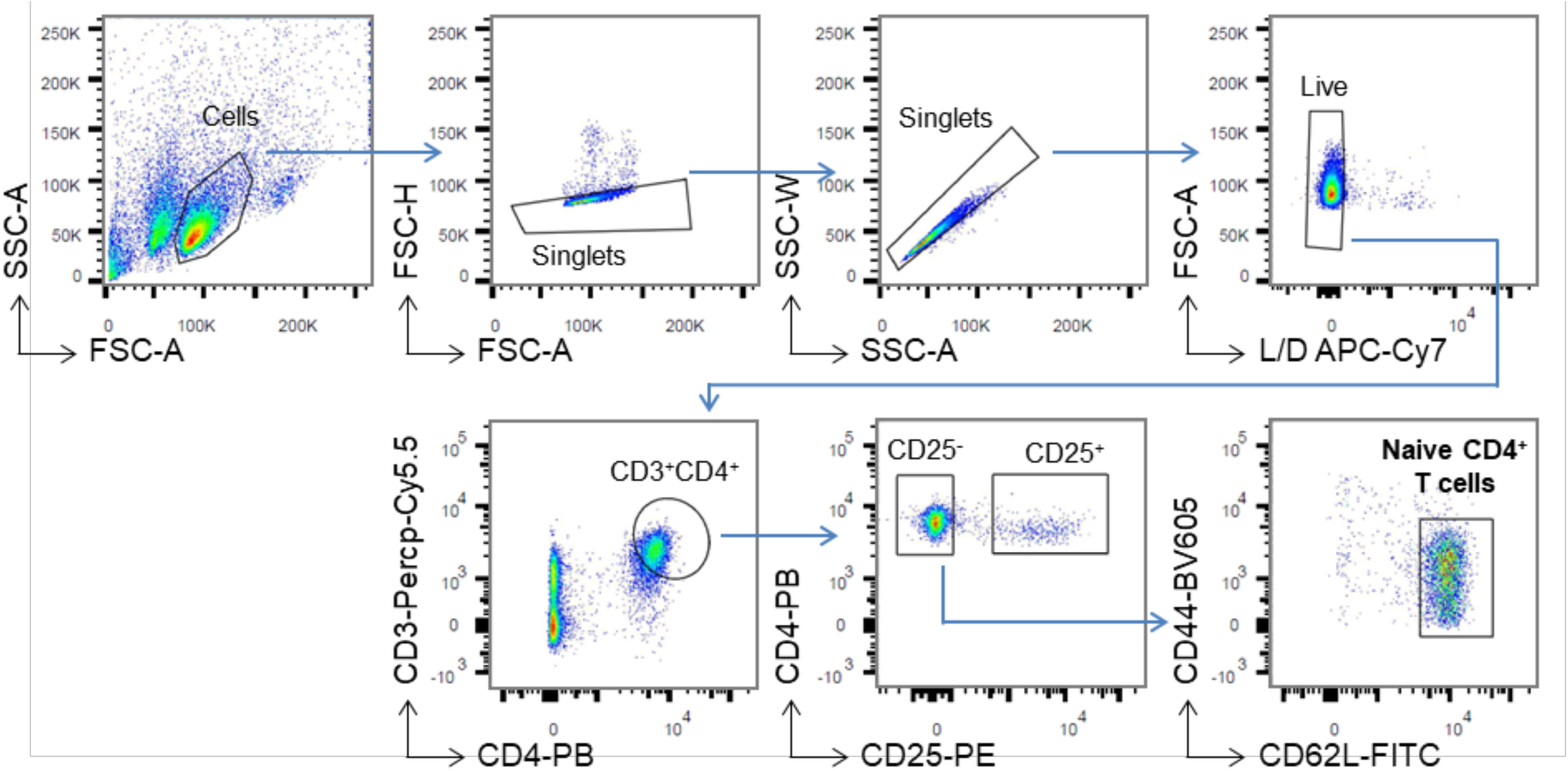
FACS gating strategy for naïve CD4^+^ T cells. Lymphocytes were selected based on a FSC/SSC plot. After selection of live singlets, lymphocytes were gated on a CD4/CD3 plot to select CD4^+^ T cells. CD4^+^ lymphocytes were then gated on a CD4/CD25 plot to exclude CD25^+^ Treg cells. Finally, CD4^+^CD25^-^ cells were gated on a CD44/CD62L to sort naïve CD4^+^ T cells.

**Fig. S3.**
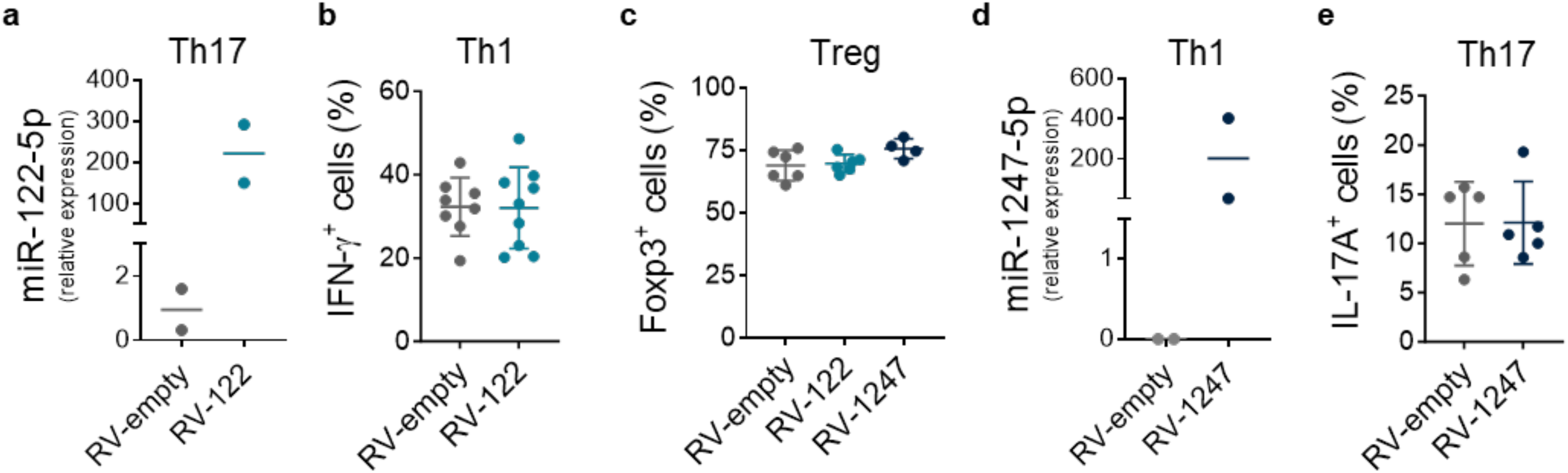
miR-122-5p and miR-1247-5p overexpression does not impact Teff and Treg differentiation *in vitro*. (a) RT-qPCR analysis of miR-122-5p expression in retrovirally transduced Th17 cells expressing a control vector (RV-empty) or miR-122 (RV-122). Results are mean from 2 independent experiments. Each dot represents cells from an individual mouse. (b) Frequency of IFN-γ^+^ CD4^+^ T cells in retrovirally transduced Th1 cells expressing a control vector (RV-empty) or miR-122 (RV-122). Results are mean ± SD from 5 independent experiments (n= 8-9). Each dot represents cells from an individual mouse. (c) Frequency of Foxp3^+^ CD4^+^ T cells in retrovirally transduced Treg cells expressing a control vector (RV-empty), miR-122 (RV-122) or miR-1247 (RV-1247). Results are mean ± SD from 3 independent experiments (n= 4-6). Each dot represents cells from an individual mouse. (d) RT-qPCR analysis of miR-1247-5p expression in retrovirally transduced Th1 cells expressing a control vector (RV-empty) or miR-1247 (RV-1247). Results are mean from 2 independent experiments. Each dot represents cells from an individual mouse. (e) Frequency of IL-17A^+^ CD4^+^ T cells in retrovirally transduced Th17 cells expressing a control vector (RV-empty) or miR-1247 (RV-1247). Results are mean ± SD from 3 independent experiments (n=5). Each dot represents cells from an individual mouse.

**Fig. S4.**
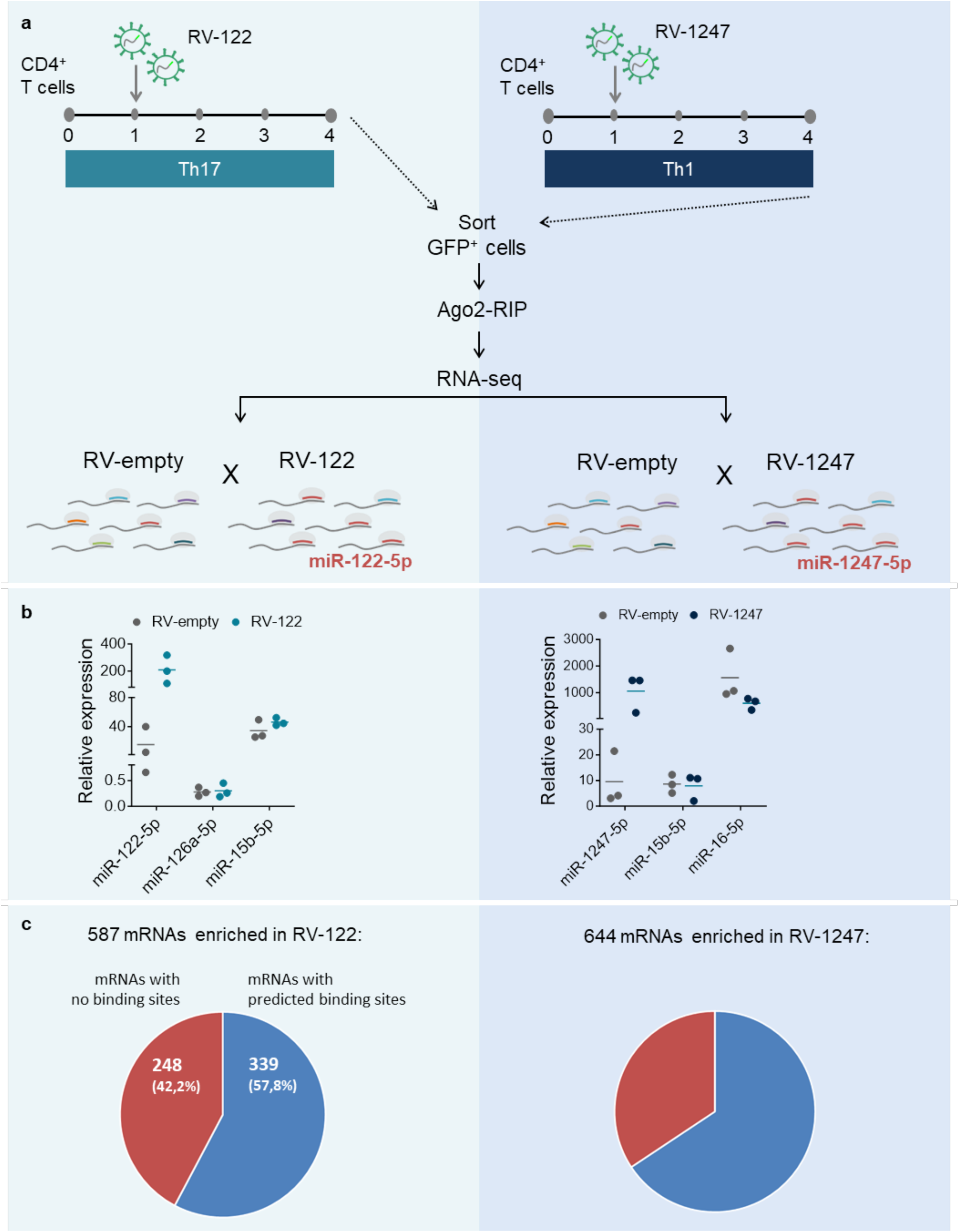
Identification of miR-122-5p and miR-1247-5p targets using differential Ago-RIPseq. (a) Experimental setup: naive CD4^+^ T cells were sorted from WT mice and differentiated *in vitro* towards Th17 or Th1 cells for 4 days. At day 1, cells were transduced with retroviral vectors encoding the precursor forms of miR-122 (for Th17) or miR-1247 (Th1) and at day 4 GFP^+^ cell were sorted. Empty vector was used as control. GFP^+^ cells were UV-crosslinked and subjected to Ago2-RNA immuniprecipitation followed by RNA sequencing to identify targets mRNA that were enriched in the samples overexpressing candidate miRNAs – miR-122 for Th17 cells and miR-1247 for Th1 cells. (b) RT-qPCR analysis of miR-122-5p, miR-126a-5p and miR-15b-5p expression after differential Ago2-immunoprecipitation of retrovirally transduced Th17 cells expressing a control vector (RV-empty) or miR-122 (RV-122) (*left panel*). RT-qPCR analysis of miR-1247-5p, miR-15b-5p and miR-16-5p expression after differential Ago2-immunoprecipitation of retrovirally transduced Th1 cells expressing a control vector (RV-empty) or miR-1247 (RV-1247) (*right panel*). Results are presented relative to miR-423-3p expression and represent mean (n=3). Each dot represents one biological replicate. (c) Following differential expression analysis, we identified 587 mRNAs overexpressed in RV-122 samples of which 57,8% (339 mRNAs) had predicted miR-122-5p binding sites (*left panels*); and 644 mRNAs overexpressed in RV-1247 samples of which 65,7% (423 mRNAs) had predicted miR-1247-5p binding sites (*right panels*).

**Fig. S5.**
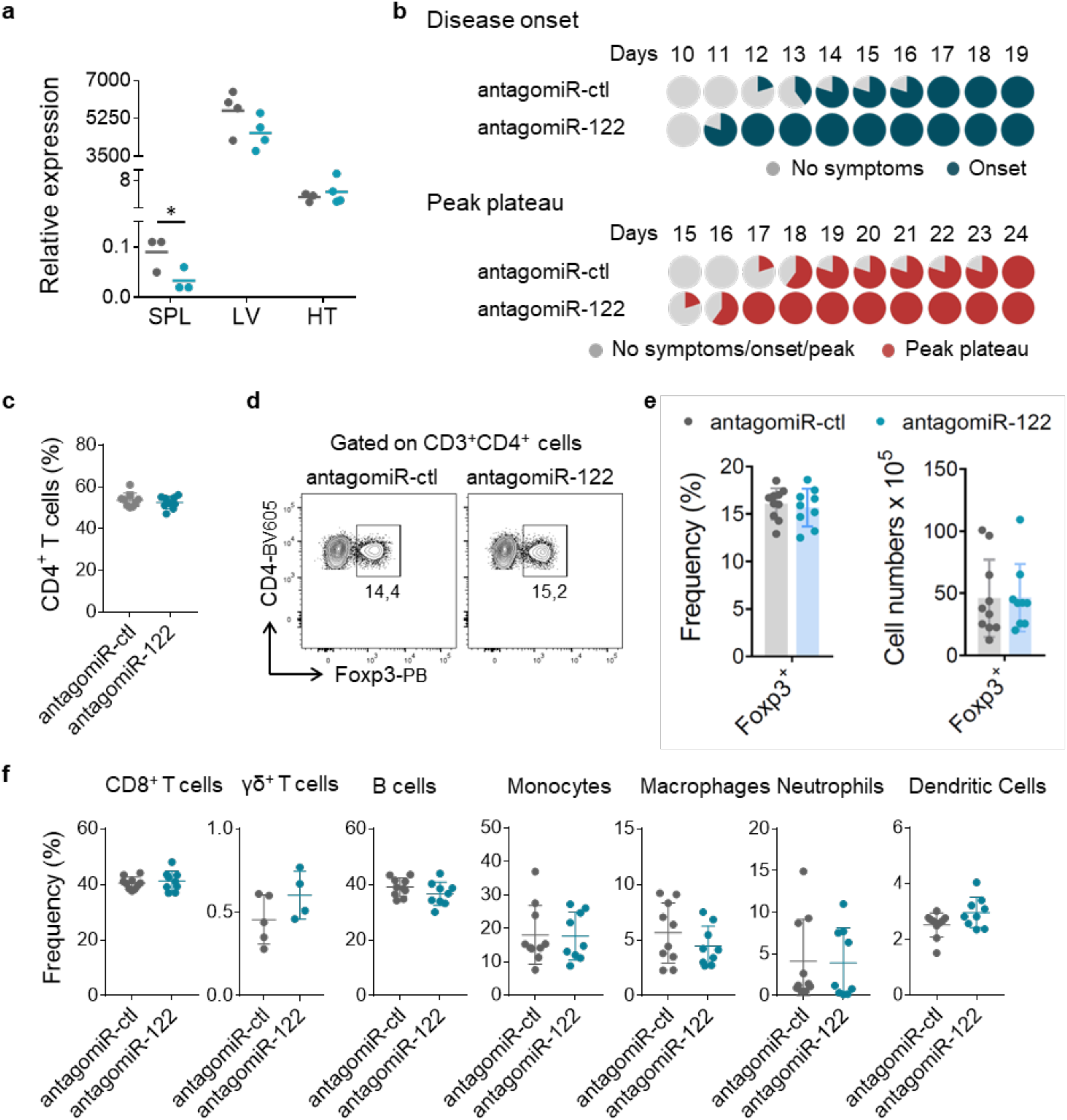
Lack of effect of miR-122-5p antagomiR in non-CD4^+^ T-cell immune cell populations. (a) RT-qPCR analysis of miR-122-5p expression in spleen (SPL), liver (LV) and heart (HT) harvested from antagomiR-treated mice at day 13 post immunization. (b) Pie charts depicting the relative number of mice treated with control and miR-122-5p antagomiRs reaching the onset of clinical symptoms or the peak plateau stage (defined as 3 days with the same score ± 0,5) in a certain day. (c) Frequency of CD4^+^ T cells in cervical and lumbar lymph nodes (cLNs) of triple Foxp3-hCD2.Ifng-YFP.Il17a-GFP reporter mice with EAE and treated with control and miR-122-5p antagomiRs. Results are mean ± SD from 3 independent experiments (n = 9-10). Each dot represents an individual mouse. (d) Representative flow cytometry plots of Foxp3^+^ CD4^+^ T cells in the cervical and lumbar lymph nodes of triple *Foxp3-hCD2.Ifng-YFP.Il17a-GFP* reporter mice with EAE and treated with control and miR-122-5p antagomiRs. (e) Frequency and absolute numbers of Foxp3^+^ CD4^+^ T cells in the cervical and lumbar lymph nodes of triple *Foxp3-hCD2.Ifng-YFP.Il17a-GFP* reporter mice with EAE and treated with control and miR-122-5p antagomiRs. Results are mean + SD from 3 independent experiments (n = 9-11). Each dot represents an individual mouse. (f) Frequency of CD8^+^ and γδ^+^ T cells, B cells, monocytes, macrophages, neutrophils and dendritic cells in cervical and lumbar LNs (cLNs) of triple Foxp3-hCD2.Ifng-YFP.Il17a-GFP reporter mice with EAE and treated with control and miR-122-5p antagomiRs. Results are mean ± SD from 3 independent experiments (n = 9-10). Each dot represents an individual mouse.

**Fig. S6.**
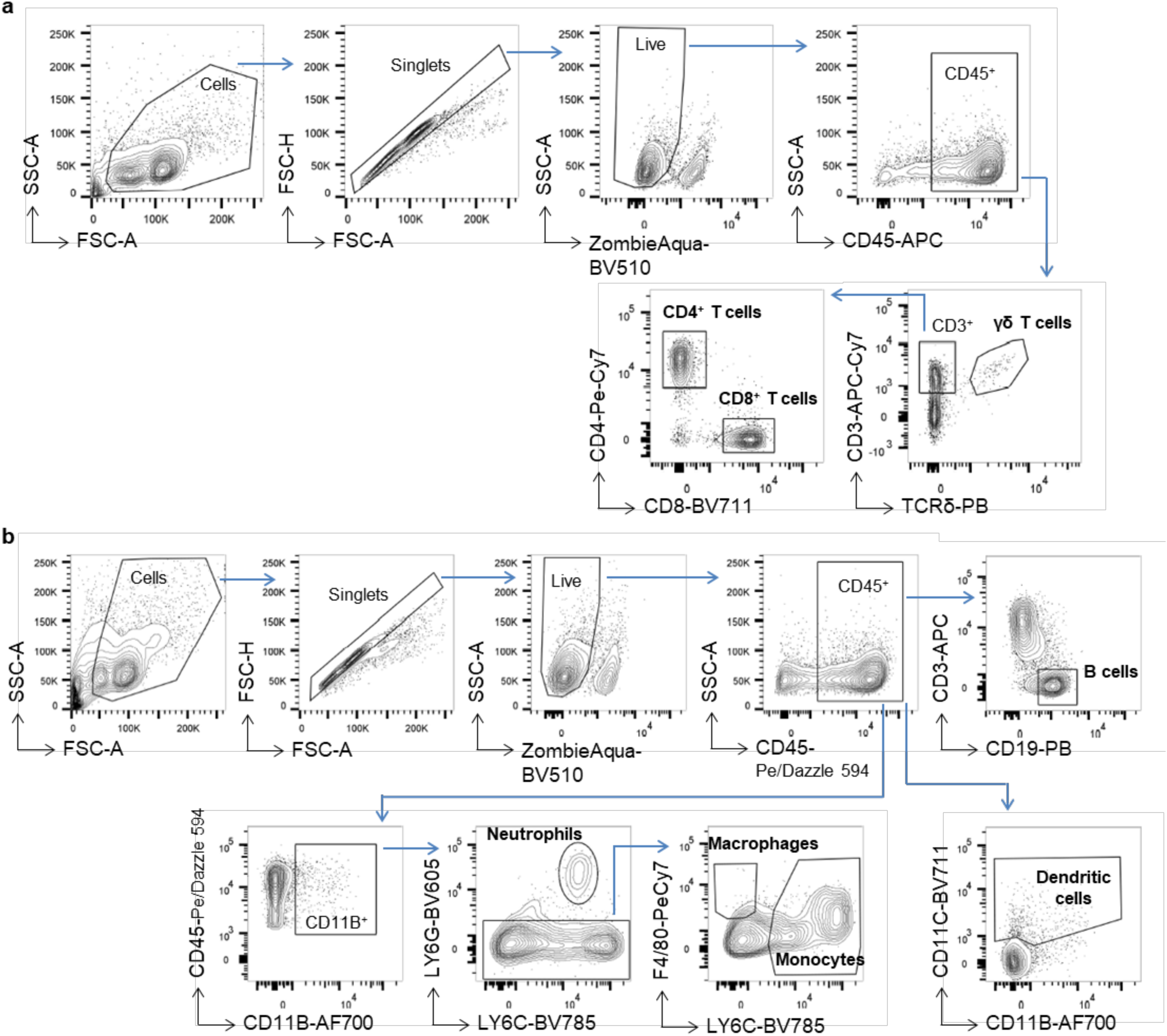
Gating strategies for immune cells populations. (a) Gating strategy for T cells: cells were selected based on a FSC/SSC plot. After selection of live singlets, lymphocytes were gated on a FSC-A/CD45 plot to select CD45^+^ immune cells. After this, lymphocyte populations were defined as following: γδ T cells - CD3^+^TCRδ^+^, CD4^+^ T cells - CD3^+^ TCRδ^-^CD4^+^CD8^-^, and CD8^+^ T cells - CD3^+^ TCRδ^-^CD4^-^CD8^+^. (b) Gating strategy for myeloid and B cells: cells were selected based on a FSC/SSC plot. After selection of live singlets, lymphocytes were gated on a FSC-A/CD45 plot to select CD45^+^ immune cells. After this, immune cell populations were defined as: B cells - CD19^+^CD3^-^, monocytes: CD11B^+^LY6G^-^LY6C^+^, macrophages - CD11B^+^LY6G^-^LY6C^-^F4/80^+^, Neutrophils - CD11B^+^LY6C^-^LY6G^+^, and dendritic cells - CD45^+^CD11C^+^.

**Fig. S7.**
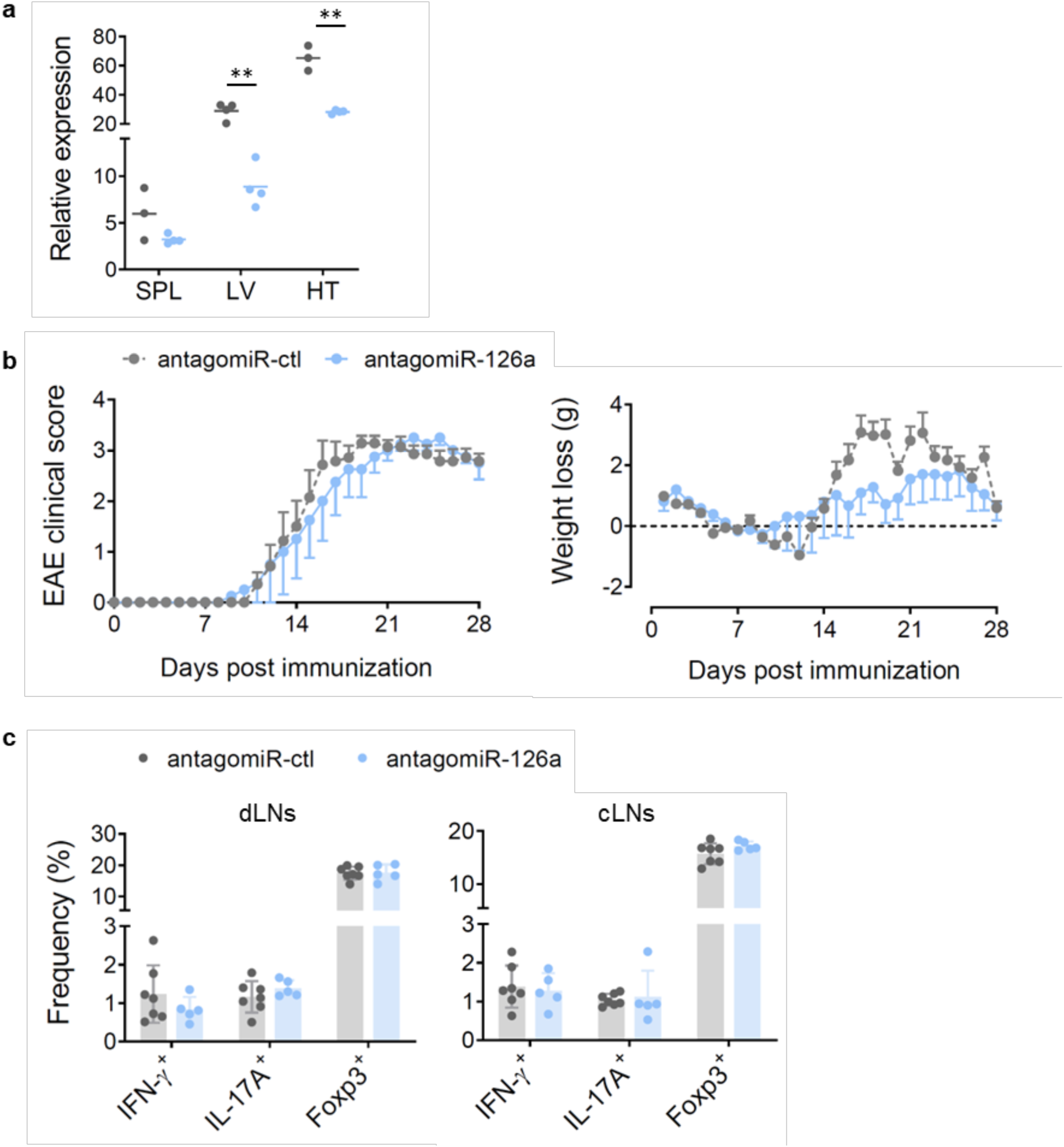
AntagomiR treatment for miR-126a does not impact EAE progression. **(a)** RT-qPCR analysis of miR-126a-5p expression in spleen (SPL), liver (LV) and heart (HT) harvested from antagomiR-treated mice at day 13 post immunization. (b) EAE clinical signs and weight daily for mice treated with miR- 126a-5p or control antagomiRs. Results are mean ± SD from 1 or 2 independent experiments (n = 3-7). Control mice are from the same experiment as shown in Figures 5 and 6. (c) Frequency of FoxP3^+^, IFN-γ^+^ and IL-17A^+^ CD4^+^ T cells in draining lymph nodes (dLNs - inguinal, axillary and brachial) and cervical and lumbar LNs (cLNs) of triple Foxp3-hCD2.Ifng-YFP.Il17a-GFP reporter mice with EAE and treated with control and miR-126a-5p antagomiRs. Results are mean ± SD from 1 or 2 independent experiments (n = 3-7). Each dot represents an individual mouse. Control mice are the same as in figure 5 and 6.

**Fig. S8.**
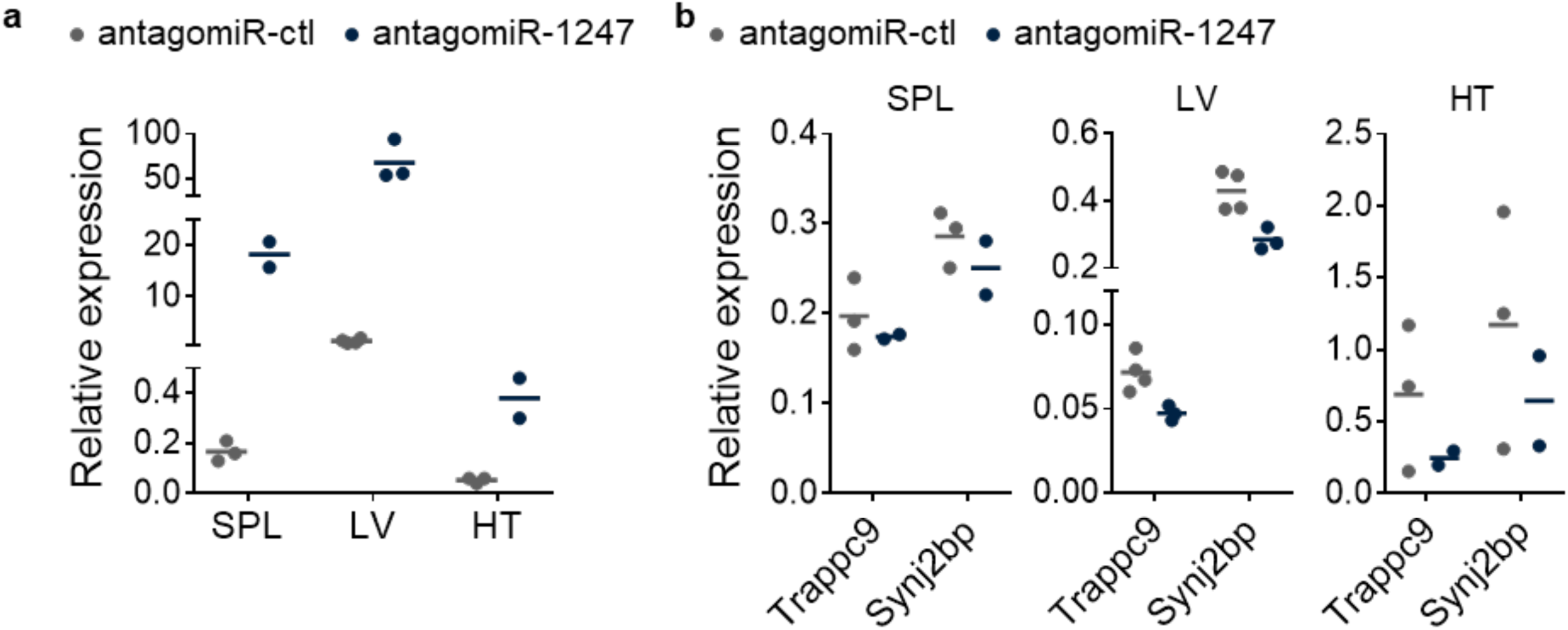
Treatment with miR-1247-5p antagomiR induces an unantecipated upregulation of miR-1247-5p levels. (a) RT-qPCR analysis of miR-1247-5p expression in spleen (SPL), liver (LV) and heart (HT) harvested from antagomiR-treated mice at day 13 post immunization. (b) RT-qPCR analysis of Trapc9 and Synj2bp expression in spleen (SPL), liver (LV) and heart (HT) harvested from antagomiR-treated mice at day 13 post immunization.

**Fig. S9.**
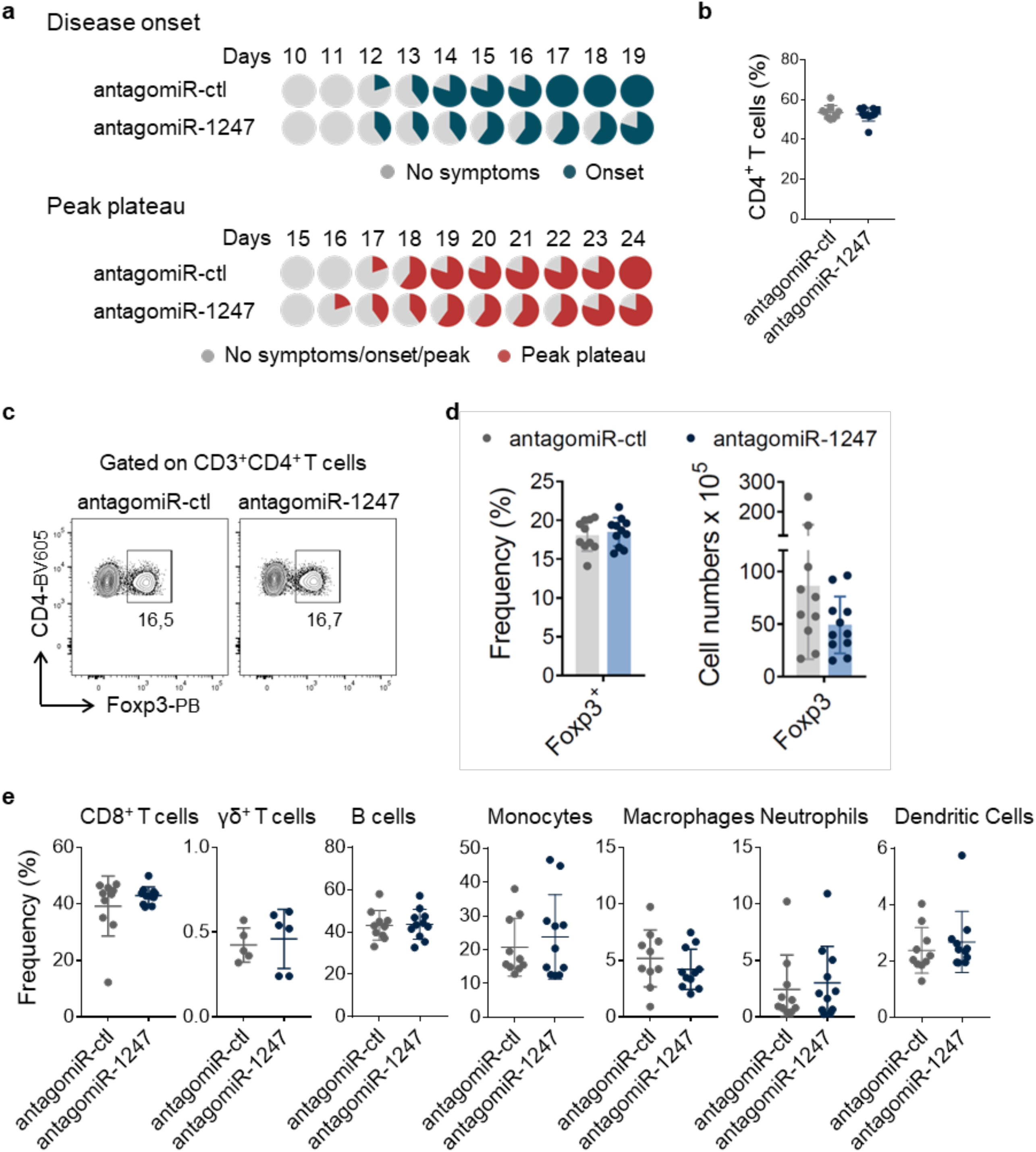
Lack of effect of miR-1247 antagomiR in non-CD4^+^ T-cell immune cell populations. (a) Pie charts depicting the relative number of mice treated with control and miR-122-5p antagomiRs reaching the onset of clinical symptoms or the peak plateau stage (defined as 3 days with the same score ± 0,5) in a certain day.(b) Frequency of CD4^+^ T cells in cervical and lumbar lymph nodes (cLNs) of triple Foxp3-hCD2.Ifng-YFP.Il17a-GFP reporter mice with EAE and treated with control and miR-122-5p antagomiRs. Results are mean ± SD from 3 independent experiments (n = 9-10). Each dot represents an individual mouse. (c) Representative flow cytometry plots of Foxp3^+^ CD4^+^ T cells in the draining lymph nodes (inguinal, axillary and brachial) of triple Foxp3-hCD2.Ifng-YFP.Il17a-GFP reporter mice with EAE and treated with control and miR-1247-5p antagomiRs. (d) Frequency and absolute numbers of Foxp3^+^ CD4^+^ T cells in the draining lymph nodes (inguinal, axillary and brachial) of triple Foxp3-hCD2.Ifng-YFP.Il17a-GFP reporter mice with EAE and treated with control and miR-1247-5p antagomiRs. Results are mean + SD from 3 independent experiments (n = 10-11). Each dot represents an individual mouse. (e) Frequency of CD8^+^ and γδ^+^ T cells, B cells, monocytes, macrophages, neutrophils and dendritic cells in cervical and lumbar LNs (cLNs) of triple Foxp3-hCD2.Ifng-YFP.Il17a-GFP reporter mice with EAE and treated with control and miR-122-5p antagomiRs. Results are mean ± SD from 3 independent experiments (n = 9-10). Each dot represents an individual mouse.

**Table S1.**
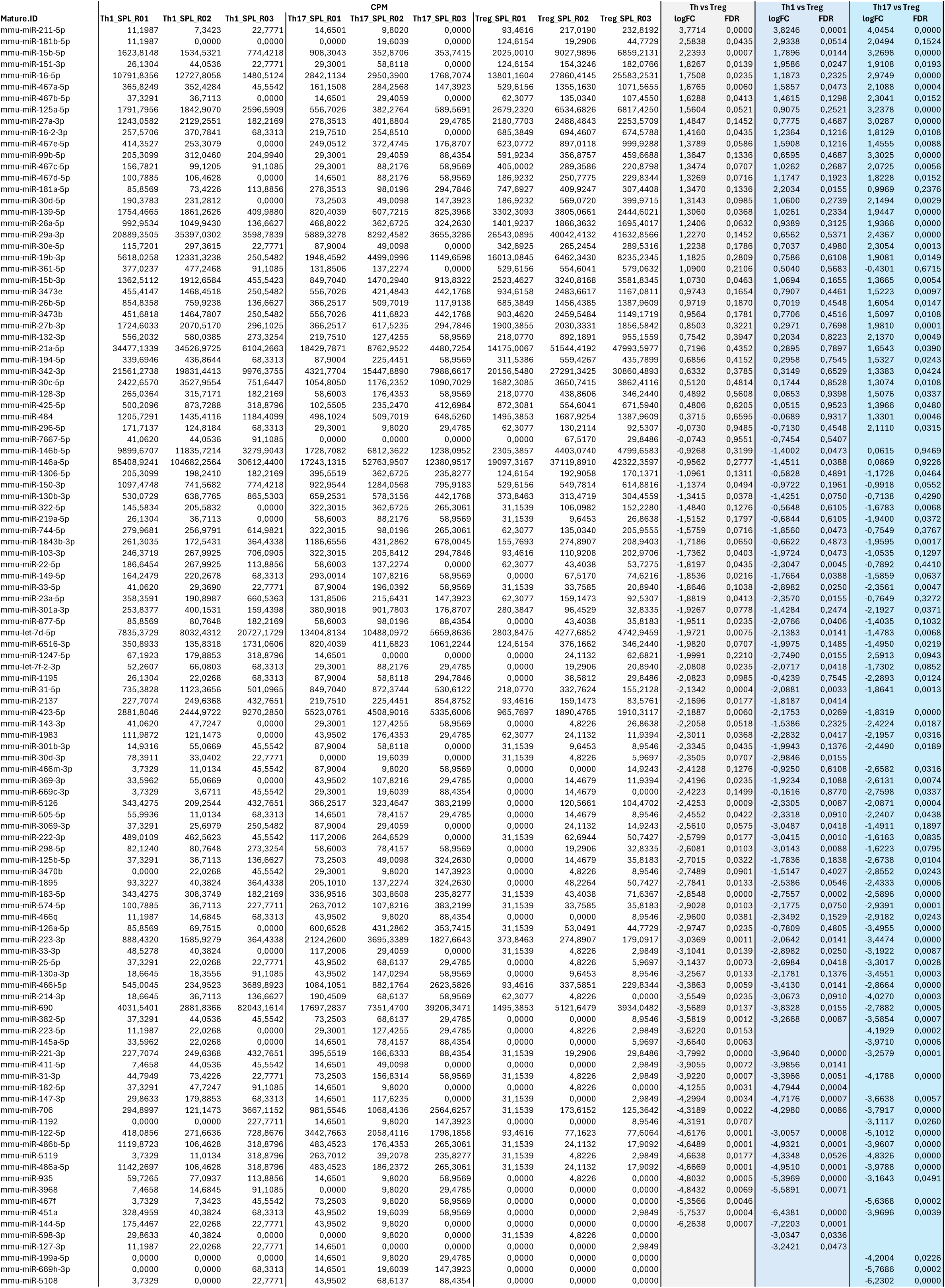
Genes differentially expressed between Teff and Treg cells.

**Table S2.**
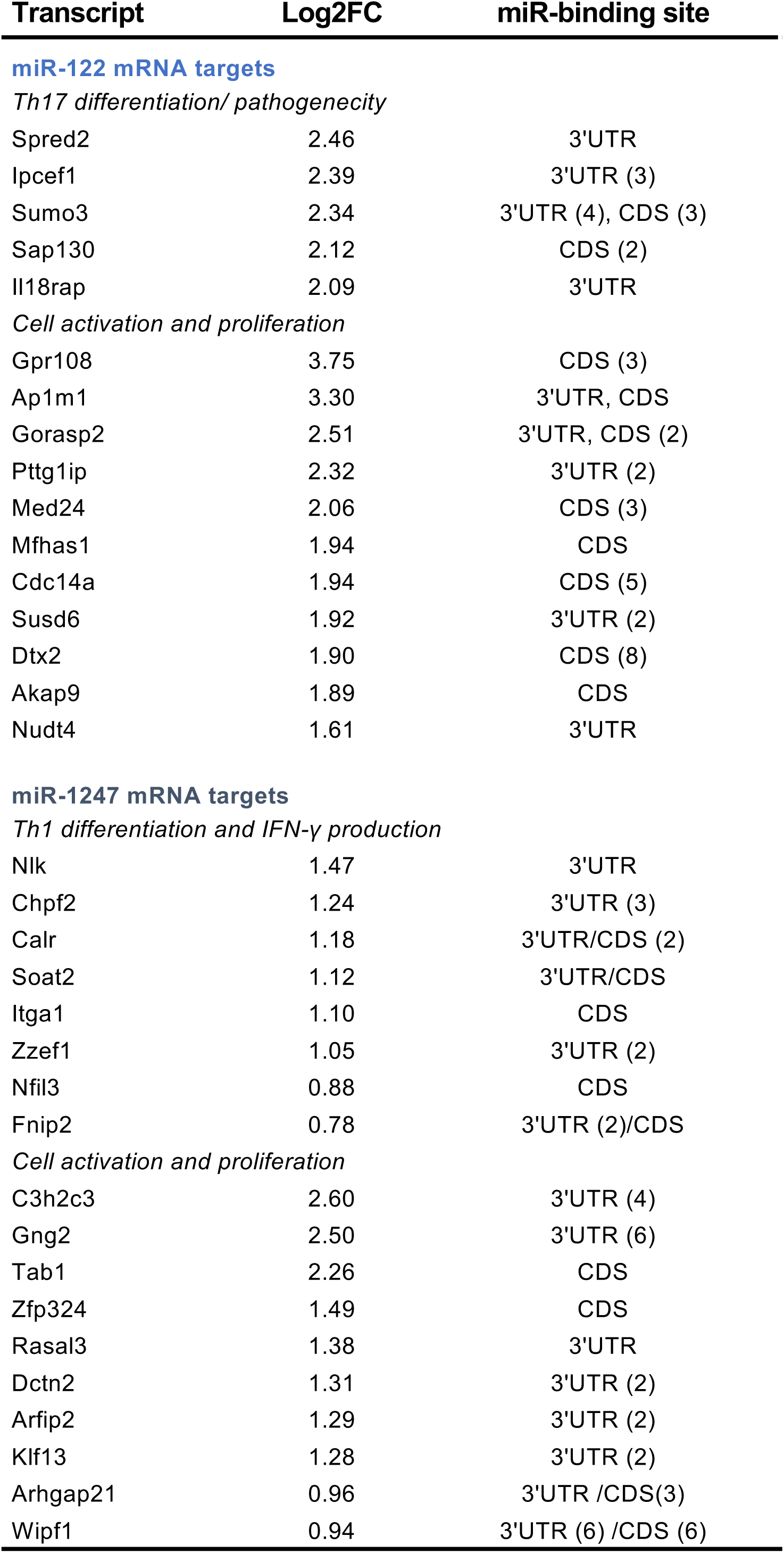
Candidate mRNA targets identified by differential Ago2-RIPseq.

**Table S3.**
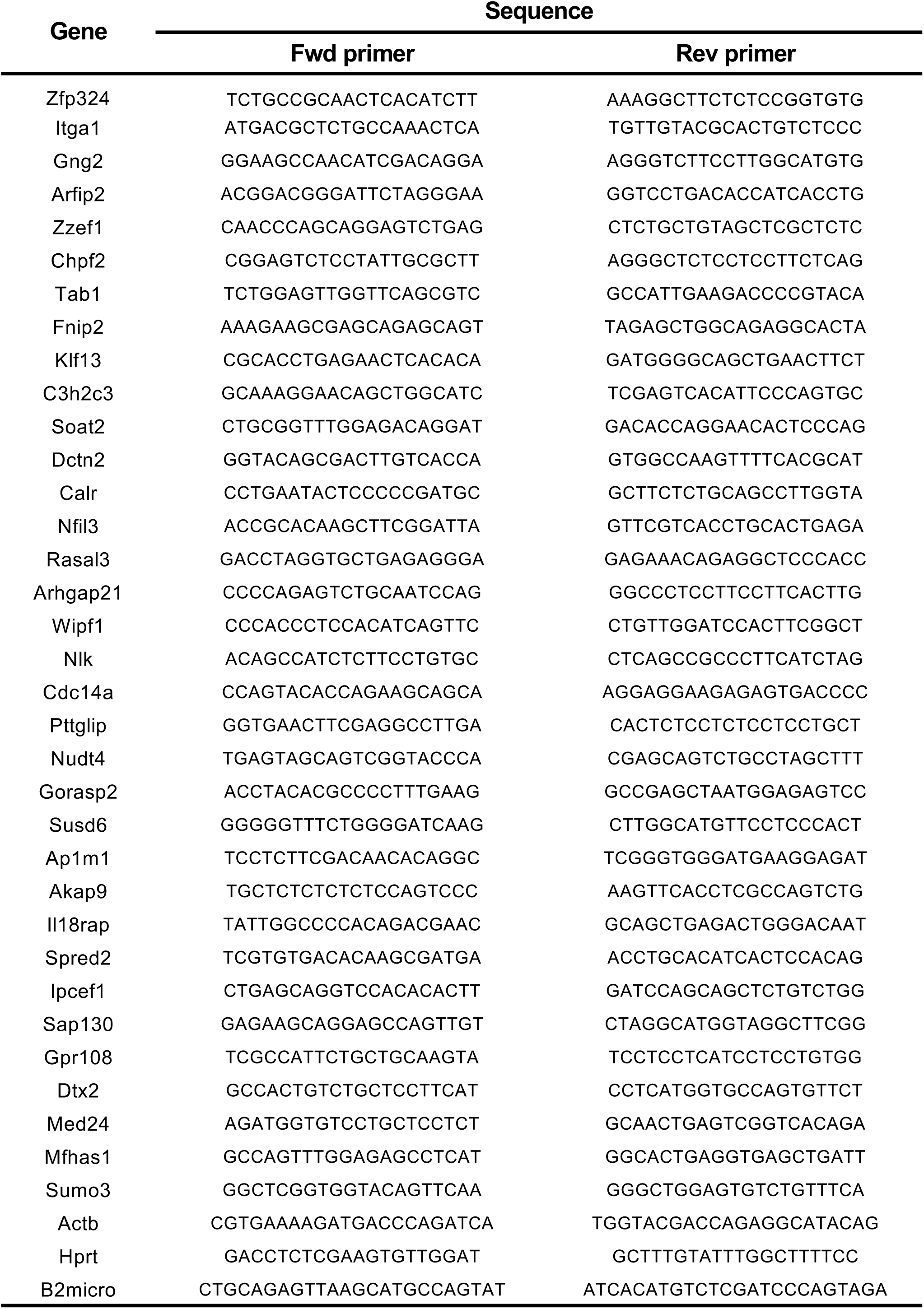
Primers used for qPCR analysis.

